# Mapping the temporal transcriptional landscape of human white and brown adipogenesis using single-nuclei RNA-seq

**DOI:** 10.1101/2022.05.30.494007

**Authors:** Anushka Gupta, Farnaz Shamsi, Mary Elizabeth Patti, Yu-Hua Tseng, Aaron Streets

## Abstract

Adipogenesis is key to maintaining organism-wide energy balance and healthy metabolic phenotype, making it critical to thoroughly comprehend its molecular regulation in humans. By single-nuclei RNA-sequencing (snRNA-seq) of over 20,000 differentiating white and brown preadipocytes, we constructed a high-resolution temporal transcriptional landscape of human white and brown adipogenesis. White and brown preadipocytes were isolated from a single individual’s neck region, thereby eliminating inter-subject variability across two distinct lineages. These preadipocytes were also immortalized to allow for controlled, *in vitro* differentiation, allowing sampling of distinct cellular states across the spectrum of adipogenic progression. Pseudotemporal cellular ordering revealed the dynamics of ECM remodeling during early adipogenesis, and lipogenic/thermogenic response during late white/brown adipogenesis. Comparison with adipogenic regulation in murine models revealed several targets for potential adipogenic/thermogenic drivers in humans. Key adipogenic and lipogenic markers revealed in our analysis were applied to analyze publicly available scRNA-seq datasets; these confirmed unique cell maturation features in recently discovered murine preadipocytes, and revealed inhibition of adipogenic expansion in humans with obesity. Overall, our study presents a comprehensive molecular description of both white and brown adipogenesis in humans and provides an important resource for future studies of adipose tissue development and function in both health and metabolic disease state.

## INTRODUCTION

Adipogenesis is a highly orchestrated process in which differentiation of adipose precursor cells (preadipocytes) into mature adipocytes is induced in response to varying metabolic needs such as energy storage during nutrient excess, lipolysis during periods of caloric deficit, or energy expenditure during cold exposure. Thus, adipogenesis is a critical process for maintaining metabolic homeostasis on an organism-wide level, and its dysregulation can contribute to diseases such as obesity, type 2 diabetes, and lipodystrophy. Consequently, there is a need to comprehensively understand the molecular regulation of adipogenic expansion in both health and in the setting of metabolic disease risk.

Adipogenic differentiation is regulated by a network of transcription factors (TFs) and results in the development of distinct types of fat from two distinct kinds of preadipocytes: white adipocytes for energy storage and brown adipocytes for thermogenic energy expenditure. Previous studies of adipogenesis’ regulation have typically employed murine model systems. For example, studies employing the murine 3T3-L1 cell line led to the identification of the core adipogenic transcriptional cascade including factors such as PPARG and C/EBPs (Ma et al. 2015; Rosen and Spiegelman 2014). Further research focused on identifying core brown versus white fat-selective factors which led to the identification of drivers and effectors of thermogenic phenotypes such as PRDM16, EBFs, and PGC1A and UCP1 (Harms and Seale 2013; Shapira and Seale 2019). More recently, modern transcriptomic investigations have identified multiple auxiliary transcription factors that serve as either positive or negative regulators of adipogenic/thermogenic response in rodents (Mota de Sá et al. 2017). Overall, studies in rodent models of adipogenesis have offered significant insights into the molecular regulation of adipogenesis. However, their applicability to humans is limited because of the existing differences in the metabolism, physiology, and transcriptomic regulation of adipose tissue between the two species. For example, BAT, which is abundantly and homogeneously present in the interscapular depot in mice, was only found to be present in adult humans over the last decade (Betz and Enerbäck 2015), and its cellular composition is heterogeneous, varying with the sampling depth in a given region (Cypess et al. 2013). Furthermore, while ADRB3 is an adrenergic receptor believed to mediate murine thermogenesis, there have been conflicting observations of its role in human thermogenesis (Cero et al. 2021; Blondin et al. 2020). Additionally, recent studies also highlighted BAT metabolic functions that do not translate from the rodents to the human (Liu, Cervantes, and Liu 2017). Consequently, there has been much interest in understanding the transcriptional control of adipocyte formation in humans (Rondini and Granneman 2020; Deutsch et al. 2020). A comprehensive understanding of the transcriptional regulation that drives adipogenesis would provide insights into lineage-determining, adipogenic, and thermogenic factors in humans, which may serve as molecular targets for therapeutic stimulation of a healthy metabolic phenotype.

Recently, multiple studies have compared the transcriptomic profiles of human-derived adipose stem cells (ASCs) at multiple stages of adipogenic differentiation using bulk gene expression profiling techniques such as microarray analysis (Ross et al. 2002; Satish et al. 2015; Urs et al. 2004; Tchkonia et al. 2005, 2006; Caserta et al. 2001), bulk RNA-seq (Ehrlund et al. 2017), and RT-qPCR (Ambele et al. 2016). This has resulted in the identification of novel adipogenic TFs such as KLFs (Wu and Wang 2013), FOXs (Gerin et al. 2009), and GATAs (Tong et al. 2000). However, bulk sampling of cells at dense time-intervals during differentiation does not allow detection of heterogeneity caused by asynchronous differentiation and the possibility of multiple lineages existing within the cellular populations of interest. By contrast, single-cell RNA-sequencing (scRNA-seq) overcomes many of these challenges and can provide an unbiased transcriptomic view of complex tissues at an unprecedented resolution (Birnbaum 2018; Trapnell 2015; Han et al. 2018). Within primary adipose tissue, recent investigations utilizing single-cell level measurements have investigated the molecular dynamics of adipocyte development in mice (Sárvári et al. 2021; Burl et al. 2018). However, single cell studies of primary tissues are not able to thoroughly sample the time course of adipogenesis. In this study, we mapped the transcriptional landscape of human white and brown adipogenesis using a unique, well-controlled, *in vitro* model system (Xue et al. 2015; Kriszt et al. 2017), which enables isolation of differentiating preadipocytes at multiple well-defined stages of development. In this system, paired white and brown primary preadipocytes were isolated from the neck of a single individual. This system, therefore, allowed us to measure transcriptional dynamics within and between white and brown lineages, while controlling for inter-individual variation typically associated with transcriptomic profiling of primary human adipose tissue, such as body mass index, genotype, and gender. Preadipocytes from both lineages were isolated and then immortalized to allow for long-term *in vitro* cell-culture. Previously reported data demonstrated highly concordant molecular features between primary and immortalized preadipocytes (Gupta et al. 2021), and *in vitro* differentiated adipocytes were shown to recover gene expression profiles and function of primary human neck BAT and WAT (Xue et al. 2015).

We now report application of single-nuclei RNA-seq (snRNA-seq) using this *in vitro* model system to perform a large-scale, time-course experiment on differentiating white and brown preadipocytes. Our use of snRNA-seq was critical to reduce bias in cell recovery, because mature adipocytes are typically incompletely recovered during single cell isolation (Deutsch et al. 2020). Our previous work demonstrated that snRNA-seq was able to capture similar cellular diversity as scRNA-seq in adipose tissue (Gupta et al. 2021). In the current work, we isolated intact nuclei from differentiating white and brown preadipocytes at 5 stages of adipogenesis (Fig 1A). We defined custom, lineage-specific adipogenic gene signatures for ordering individual nuclei in increasing order of maturity. Our analyses revealed temporal regulation of distinct gene modules in both white and brown adipogenesis, each module highlighting the dynamics of biologically relevant functional processes. We further used our dataset to understand cell maturity differences across distinct metabolic phenotypes and cell-types.

**Fig. 1.**
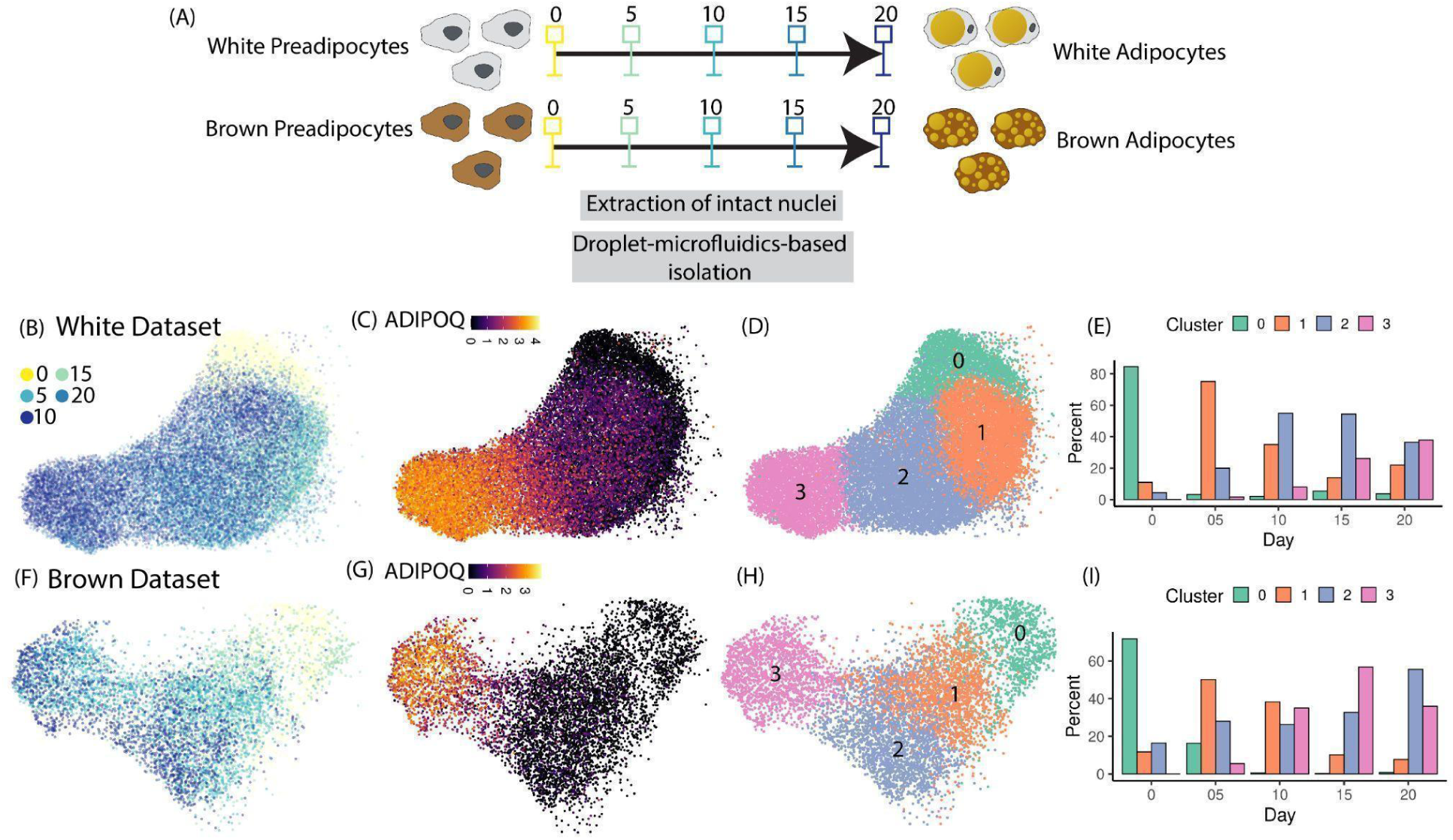
snRNA-seq of differentiating preadipocytes enables high-resolution sampling of adipogenic cellular states. **(A)** Schematic of the experimental design utilized in this study. **(B) to (D)** UMAP visualization of white adipogenesis dataset colored by (B) day of collection (C) *ADIPOQ* gene expression and (D) unsupervised cluster classification. **(E)** Distribution of nuclei harvested at each experimental time-point with clusters identified in (D). **(F) to (I)** Same plots as (B) to (E) but plotted for brown adipogenesis dataset.

## RESULTS

### snRNA-seq enables high-resolution sampling of adipogenesis

Cultured white and brown preadipocytes were differentiated into respective mature adipocyte types using an induction cocktail (see Methods). During differentiation, intact nuclei were harvested from white and brown preadipocytes (see Methods) at 5 equally spaced time-points during the 20-day adipogenic induction period (Fig. 1A). Isolated nuclei were subjected to droplet-based snRNA-seq, followed by QC analyses (see Methods). In total, we recovered 25,339 high-quality nuclei from white precursors and 27,568 high-quality -nuclei from brown precursors, with ∼2000-6000 genes detected per nucleus (Table 1).

**Table 1:** Sequencing metrics for snRNA-seq libraries analyzed in this study.

|  | White Adipogenesis |  | Brown Adipogenesis |  |
| --- | --- | --- | --- | --- |
| Day | Number of nuclei recovered | Reads per nuclei | Number of nuclei recovered | Reads per nuclei |
| 00 | 8026 | 174,738 | 6763 | 216700 |
| 05 | 39,345 | 37,939 | 9107 | 94,812 |
| 10 | 7,229 | 102,155 | 9945 | 83,732 |
| 15 | 4,834 | 174,967 | 3217 | 122,859 |
| 20 | 12,160 | 123,700 | 8085 | 181,758 |

Independent analysis of white preadipocytes across 5 time-points revealed detection of adipocytes as soon as day-5 (Fig. S1A), with continuously increasing expression for the mature adipocyte marker gene *ADIPOQ* and decreasing expression for progenitor marker *CD44* (Fig. S1A) over the time course. Integration of these 5 datasets using scVI-tools (Gayoso et al. 2021) revealed structuring of nuclei along a continuum of cellular states (Fig. 1B), starting from early precursors to mature adipocytes (as marked by *ADIPOQ*, Fig. 1C). Unsupervised clustering revealed four distinct clusters during white adipogenesis (Fig. 1D), with the majority of day-0 nuclei grouped in cluster-0 and majority of day-20 nuclei grouped in cluster-3, thereby illustrating the increasing maturity level from cluster-0 to cluster-3. Nuclei harvested at later time points were distributed over all clusters (Fig. 1E), thereby highlighting the asynchronous behavior of adipogenic differentiation in the *in vitro* model system.

For brown adipogenesis, preadipocytes showed heterogeneity at day 0 (Preadipocyte-1 and Preadipocyte-2, Fig. S1B), with the Preadipocyte-2 population expressing genes that correspond to adipogenic capacity (Gupta et al. 2021). Independent analysis of brown nuclei harvested at later time-points (day-5 to day-20) showed similar preadipocyte heterogeneity along with capture and detection of mature adipocytes (Fig. S1B). Integrative analysis of all 5 datasets confirmed that Preadipocyte-2 population differentiates into mature adipocytes, while Preadipocyte-1 appears to exhibit a non-adipogenic, non-thermogenic response pattern (Fig S1C). Notably, pathway analysis using genes up regulated as part of the non-adipogenic response revealed enrichment of FOXO1 transcriptional activity (Fig. S1D), a known repressor of PPARG (Chen et al. 2019; Fan et al. 2009), suggesting a possible role of FOXO1 in inhibiting adipogenic response in the Preadipocyte-1 population.

In the brown adipogenic cluster, nuclei harvested across different days were distributed over a continuum of maturation state (Fig. 1F), with a continuously increasing expression of *ADIPOQ* (Fig. 1G). Like white adipogenic response, unsupervised clustering identified 4 clusters during brown adipogenic response (Fig. 1H), each with increasing maturity level (Fig. 1I). Of note, nuclei harvested on day-20 were primarily grouped in cluster 2 whereas nuclei harvested on day-15 were primarily grouped in cluster 3 (Fig. 1I). We attributed this observation to a lower differentiation efficiency on day-20, thereby resulting in reduced number of harvested mature adipocytes (grouped in cluster 3) as compared to day-15. We performed adipogenic signature analysis, in which a score is assigned to each nucleus based on expression of adipogenic genes, that revealed the highest score for day-20 adipocytes in cluster-3 (Fig. S1E). As compared to white adipocytes, differentiated brown adipocytes had up-regulation of the brown-adipocyte-specific marker gene *ZIC1* (Sharp et al. 2012) *as well as PGC1B*, a regulator of hepatic glucose and lipid metabolism (Medina-Gomez, Gray, and Vidal-Puig 2007) and a recognized thermogenic marker (Hussain, Roesler, and Kazak 2020); Fig. S1F). Analysis of differential expression followed by transcription factor enrichment analysis between mature white and brown adipocytes (cluster-3 in Fig. 1D vs cluster-3 in Fig. 1H) identified that the top-ranking transcription factor enriched in brown adipocytes was *FOXS1* (Heglind et al. 2005) *and FOXC2* (Cederberg et al. 2001). Thus, our experimental strategy allowed us to capture a spectrum of cell-states undergoing differentiation toward white or brown adipocyte lineages.

### Pseudotemporal ordering of differentiating preadipocytes identifies dynamics of extracellular matrix (ECM) remodeling, lipogenesis, and thermogenesis

#### Lineage-specific gene signatures enable high-resolution ordering of single nuclei

Dense sampling of cellular states with snRNA-seq enabled reconstruction of the adipogenic developmental trajectory by ordering individual nuclei along a pseudo-time axis. To achieve this, we identified a set of genes specific to both white and brown-adipogenesis that defined progression through differentiation and provided a signature score that was used as a proxy for pseudo-time (Note S1). These gene signatures consisted of genes that showed monotonic increase in expression from immature preadipocytes to mature adipocytes (Note S1). Hence, pseudo-temporal scoring based on expression of such genes provided a high dynamic range and resolution of cellular maturation state assignment.

Using the white-/brown-adipogenesis-specific gene signature, differentiating preadipocytes were ordered by increasing maturation state (Fig. 2A and 2C). Assessment of expression dynamics in pseudotime of key adipogenic TFs *CEBPB, CEBPD, PPARG* and *CEBPA* accurately reflected the core adipogenic signaling cascade (Fig. 2B and 2D), with early induction of *CEBPB*, followed by sustained expression in response to insulin (see Methods, (MacDougald et al. 1995), early induction and transient expression of *CEBPD*, stable increase in expression of *PPARG*, and late induction of *CEBPA*, thereby validating our cell-ordering strategy for both white and brown adipocyte development. Next, dynamically regulated genes were identified by binning nuclei in the pseudotemporal space (Fig. S2A and S2B, see Methods, Steier et al., 2021). We reasoned that there would be significant deviation in expression of dynamically regulated genes from either the initial or final stages of adipogenesis, and hence performed differential expression testing for each bin against the first and last pseudotemporal bins (logFC >1 and FDR < 0.05) to identify such genes. In total, we identified 596 and 454 temporally expressed genes during white and brown adipogenesis respectively. Unsupervised hierarchical gene clustering identified three major expression trends (Fig. 2E to 2H, Table S1 and S2) which we describe as down-regulated (Module 1 and Module 2), transiently up-regulated (Module 3), and up-regulated (Module 4 and Module 5). Within down-regulated gene modules, Module 1 undergoes immediate down-regulation, whereas Module 4 undergoes a consistent down-regulation (Fig. S2I and S2J). Within up-regulated gene modules, Module 4 has a consistent up-regulation whereas Module 5 has a delayed up-regulation (Fig. S2K and S2L).

**Fig. 2.**
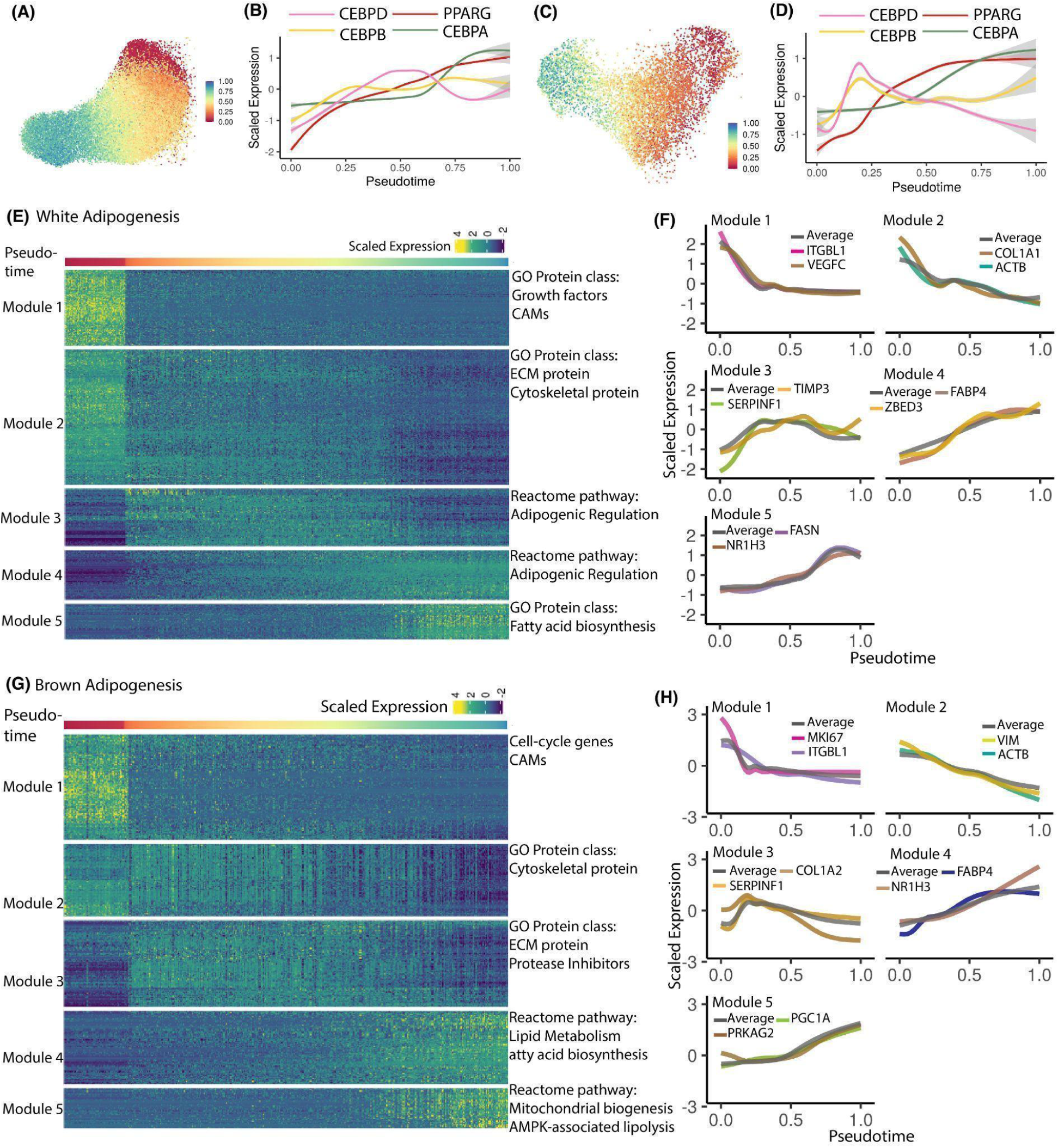
Gene module identification reveals expression dynamics of key biological processes accompanying white and brown adipogenic progression. **(A)** Pseudotemporal ordering of differentiating white preadipocytes. **(B)** Expression dynamics for core adipogenic transcription factors CEBPs and PPARG during white adipogenesis. **(C) and (D)** Same plot as (A) and (B) but for brown adipogenesis. **(E)** Heatmap of expression dynamics for gene modules identified in white adipogenesis. Each column indicates a single nucleus, with columns ordered by increasing pseudotemporal score. Each row depicts a single gene, with rows clustered by unsupervised Louvain clustering. Each gene module is annotated by terms reflecting key biological processes based on Gene Ontology. **(F)** Smoothed expression dynamics of selected genes from each gene module identified in panel (E) in white adipogenesis dataset. **(G) and (H)** Same plot as (E) and (F) but for brown adipogenesis dataset

#### Cell-adhesion disruption is followed by fibrillar to basement-membrane-type ECM remodeling during early adipogenic progression

ECM remodeling during adipogenesis is key to providing the appropriate niche as spindle-shaped preadipocytes transform into spherical, fragile, lipid-laden adipocytes (Mariman and Wang 2010; Nakajima et al. 2002). Dysregulation of ECM remodeling is a hallmark of clinical obesity, in which adipose tissue becomes fibrotic because of increased deposition of fibrillar ECM components such as collagen type 1, -3, and -5 (DeBari and Abbott 2020). While ECM remodeling is central to healthy adipogenic expansion, the precise dynamics of ECM reorganization and its molecular regulation remains poorly understood. Here, we focused on the differential dynamics across Modules 1, 2, and 3 (Fig. S2I and S2J) to provide novel insights into the temporal regulation of ECM reorganization during human adipogenic progression.

During white adipogenesis, genes undergoing rapid down-regulation upon addition of adipogenic medium (Module 1) primarily included cell adhesion molecules (CAMs) such as *ITGB8, ITGA11*, and *ITGBL1*, as well as growth factors such as *VEGFA, VEGFC, FGF2*, and *FGF5* (Fig. 2F, Table S2 and S4). These observations are supported by previous studies which report downregulation of such integrin-associated genes during adipogenesis (Spurgin et al. 2016; Ullah, Sittinger, and Ringe 2013; Morandi et al. 2016), and known anti-adipogenic effects of these growth factors (Kim et al. 2015; Côté et al. 2017; Karaman et al. 2016). Notably, CAMs serve as contact points between cells and ECM; thus, down-regulation of CAMs suggests that disruption of cellular-ECM contacts is a very early response to the induction media. Module 2 genes undergoing more gradual down-regulation primarily included ECM structural components (Fig. 2F, Table S2 and S4). such as fibrillar collagen types *COL1, COL3*, and *COL5*. This observation agrees with previous reports and suggests that degradation of such ECM components paves the way for basement-membrane-type ECM components such as collagen-4 which are better suited to support spherical adipocytes (Mor-Yossef Moldovan et al. 2019; Al Hasan et al. 2021). Indeed, expression of collagen-4 gradually increased during white adipogenesis in our dataset (Fig. S2C, Table S2). Module 2 also included down-regulated cytoskeletal components such as actin (*ACTB)*, tubulin *(TUBB)*, and vimentin *(VIM)* (Fig. 2F, Table S2 and S4), in agreement with previous reports (Spiegelman and Farmer 1982; Yang et al. 2013). Module 3 genes (transient up-regulation) mostly consisted of protease inhibitors such as *TIMP3* and *SERPINF1* (Fig. 2F, Table S2 and S4). Protease inhibitors serve as ECM constructive enzymes, antagonizing the ECM degradation activity of metalloproteases such as MMPs, ADAMs, and ADAMTSs (Lilla et al. 2002). Notably, all such metalloproteases were downregulated during white adipogenesis in our dataset (Fig. S2D). Therefore, our results are consistent with an initial disruption of cell adhesion contacts, followed by metalloprotease activity to promote ECM degradation and a subsequent shift toward ECM regeneration via activity of protease inhibitors.

Brown adipogenesis recovered similar dynamics of ECM remodeling, with immediate downregulation of integrins and CAMs such as *ITGBL1* (Fig. 2H), *CDH11*, CDH13, *and CDH2* (Table S1). Similarly, Module 2 included gradually downregulated cytoskeletal components such as *ACTB, TUBB*, and *VIM* (Fig. 2H, Table S1 and S5). However, unlike white adipogenesis, fibrillar collagen components such as collagen types -1, -3, and -5 were clustered in Module-3 with initial up-regulation (Fig. 2H, Table S1 and S5), likely providing a fibrillary-type ECM to support early proliferation of brown preadipocytes (Fig. S2E and S2F) until their growth arrest (Mor-Yossef Moldovan et al. 2019). Finally, several patterns were very similar to those observed during white adipogenesis. Collagen types -1, -3, and -5 are downregulated over time, with a converse increase in expression of basement-membrane-type collagen-4 (Fig. S2G). Likewise, metalloprotease inhibitors were enriched in Module-3 (Fig. 2H, Table S5) with consistent downregulation of metalloproteases (Fig. S2H), suggesting a similar shift in enzymatic activity from ECM degeneration to ECM reconstruction.

#### Differential dynamics of adipogenic, lipogenic, and thermogenic response marks late white and brown fat development progression

Typically, fat synthesis (or lipogenesis) accompanies later stages of adipogenesis as preadipocytes accumulate lipid droplets characteristic of mature adipocytes. However, previous studies have identified distinct pathways and regulators specific to lipid droplet biogenesis, expansion, and shrinkage during fat cell maturation (Yu and Li 2017; Meex, Schrauwen, and Hesselink 2009), indicating a lipogenesis-specific transcriptional network. Similarly, brown-fat-specific transcriptional networks which regulate thermogenesis have been uncovered (Patrick Seale et al. 2008; Shapira and Seale 2019). Hence, we investigated the dynamics of adipogenic, lipogenic, and thermogenic responses in human white and brown adipogenesis dataset. Specifically, we focused on gene modules 4 and 5, which are upregulated in response to adipogenic induction, but with differential dynamics (Fig. S2K and S2L).

In white adipogenesis, Module 4 primarily consisted of genes that regulate lipid mobilization such as *PLIN1, FABP4, CD36* and adipogenic transcriptional regulators such as *PPARG, MLXIPL* (Hurtado Del Pozo et al. 2011), *and ZBED3* (Xu et al. 2021), all of which were gradually up-regulated (Fig. 2F, Table S2). Module-5, on the other hand, included lipogenic genes such as *FASN, ACSL1, GPAM* and the lipogenic transcription factor *NR1H3* (Schultz et al. 2000), Fig. 2F, Table S2), suggesting a delayed onset of lipogenic transcriptional response when compared to an adipogenic response. This agrees with pathway analysis which revealed enrichment of adipogenic regulation terms in Module-4 (Table S4) and fatty acid biosynthesis terms in Module-5 (Table S4). Therefore, our results indicate that adipogenic transcriptional responses precede lipogenic responses during white adipocyte development.

By contrast, patterns were distinct in brown adipogenesis. Module 4 genes were enriched for transcriptional regulators of both adipogenic and lipogenic response such as *PPARG, FABP4, SREBF1* (Fig. 2H, (Shao and Espenshade 2012; Talebi et al. 2018), and *NR1H3* (Fig. 2H, (Schultz et al. 2000). Pathway analysis also identified enrichment of both fatty acid biosynthesis, and lipid metabolism-associated terms in Module 4 genes (Table S5). This adipogenic and lipogenic response was followed by a thermogenic response (Table S5), as indicated by the enrichment of AMPK-associated lipolytic pathways (Herzig and Shaw 2018; Ahmadian et al. 2011) and mitochondrial biogenesis pathways (Table S5) in Module 5. Overall, our results suggest a parallel adipogenic & lipogenic response during brown fat development, followed by a later thermogenic response.

#### High-resolution map of transcription factor dynamics identifies potential regulators of adipogenic and thermogenic response in humans

Differential expression patterns of transcription factor mRNAs can help identify distinct cell-types, -states, or -lineages. Using our snRNA-seq dataset, we specifically characterized and compared the expression dynamics of transcription factors during white and brown fat development.

During white adipogenesis, we identified 49 TFs with dynamic gene expression profiles (Fig. 3A). As expected, most of these TFs had similar expression dynamics as previously reported in rodents (Fig. 3A). This included Module 1 anti-adipogenic TFs such as *GLI2* (hedgehog signaling mediator (Shi and Long 2017; Fontaine et al. 2008), *RBPJ* (Notch signaling mediator(Bi et al. 2014; Shan et al. 2017), and *AHRR (Ishihara, Tsuji, and Vogel 2018)*,Module 2 anti-adipogenic TFs such as *TCF4* & *TCF12* (mediator of Wnt/B-catenin (Hrckulak et al. 2018), and *SMAD3* (mediator of TGFB pathway4 (Choy, Skillington, and Derynck 2000), Module 4 pro-adipogenic TFs such as *PPARG, MLXIPL*, and *ZBED3*, and Module 5 pro-lipogenic TF *NR1H3*. We also identified multiple TFs with dynamic gene expression profiles during white fat development that had not previously been associated with adipogenesis in humans (Fig. 3A, highlighted in red), and which may serve as potential adipogenic regulators in humans. This included KLF12, ZEB2, CREB3L2, and MEF2A (Fig. 3A); paralogs of these genes KLF8 (Lee et al. 2012), ZEB1 (Gubelmann et al. 2014), CREB5 (Reusch, Colton, and Klemm 2000), and MEF2D (Li et al. 2015) are known regulators of adipogenesis in rodents, thereby suggesting a similar role for these genes in humans. Moreover, transcription factor binding site enrichment analysis demonstrated overrepresentation of RFX8, BNC2, and TSHZ3 binding sites in differentially expressed genes.

**Fig. 3.**
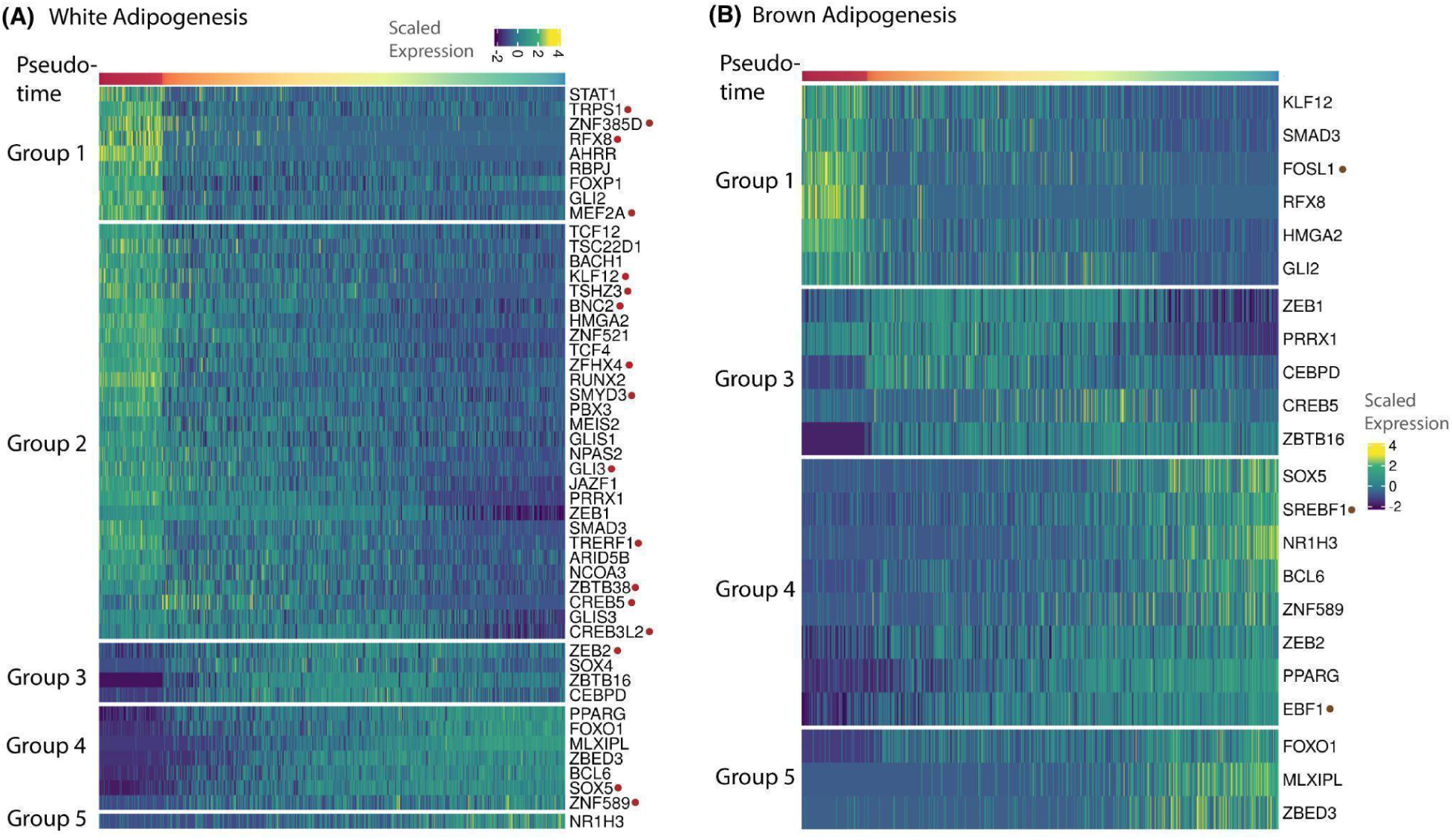
Transcription factor dynamics in white and brown adipogenesis. **(A)** Dynamics of all TFs temporally regulated during white adipogenesis, distributed by their gene module annotation. Highlighted in red are TFs that had not previously been associated with adipogenesis in humans **(B)** Dynamics of all TFs temporally regulated during brown adipogenesis, distributed by their gene module annotation. Highlighted in brown are TFs that are exclusively regulated during brown adipogenesis

While many studies have focused on drivers of adipogenesis, this data provided the opportunity to identify thermogenic TFs based on unique regulation in brown adipogenesis over white adipogenesis, in a model system derived from the same individual. Typically, thermogenic TFs are identified based on differential enrichment of genes in BAT over WAT. While such a strategy is applicable in rodents, in which BAT is predominantly localized in a discrete interscapular depot, it is challenging in humans where BAT is found interspersed within WAT. Moreover, such a strategy provides a tissue-level measurement, with no information on whether identified differences manifest at the early preadipocyte stage or late mature adipocyte stage. Our dataset mitigated this challenge by allowing us to investigate the temporal dynamics of TF expression during fat development across multiple stages of adipogenesis. In total, we identified temporal regulation of 22 TFs, a majority of which were similarly regulated during white adipogenesis (Fig. 3B and Fig. S3A). This included novel TFs RFX8, KLF12, CREB5, ZEB2, SOX5, and ZNF589 identified in the paragraph above (Fig. S3A). Notably, we identified exclusive significant regulation of TFs SREBF1, EBF1, and FOSL1 during brown adipogenesis (Fig. 3B). Of these, SREBF1 and EBF1 were minimally regulated during white adipogenesis (Fig. S3B), which agrees with their previously demonstrated pro-thermogenic roles (Shapira and Seale 2019; Sanchez-Gurmaches et al. 2018). Next, we identified TFs with exclusive activity in brown adipogenesis using binding-site enrichment analysis (Table S3). This included thermogenic regulatory TFs such as *SREBF1, KLF15* (Yamamoto et al. 2010), *and TWIST1 (P. Seale 2010)*, along with TFs *FOXL1, ZNF117, IRX6, OSR1*, and *PRRX2*, whose involvement in the context of thermogenic response has not been previously characterized. TFs *ZNF117* and *IRX6* were enriched in up-regulated genes (Module-4 and Module-5, Table S3) along with pro-thermogenic TFs *SREBF1* and *KLF15*, thereby suggesting a potentially similar function for these TFs in human thermogenesis.

#### Adipogenic gene signature reveals differences in cell maturation state within previously characterized distinct white preadipocyte types

Utilization of gene signatures in our dataset enabled the ordering of differentiating preadipocytes by their maturation state. The applicability of these signatures can also be extended to other primary adipose tissue scRNA-seq/snRNA-seq datasets to investigate differences in cell maturation state that may be driven by factors such as metabolic phenotype, diseased state, BMI etc. This can be achieved by using appropriate signatures (based on the lineage) as input to Vision (DeTomaso et al. 2019) to assign scores to cells of interest in a given dataset. The assigned score can then be tested for statistical difference, thereby providing comparative, quantitative insights into maturation state differences. Since scores assigned using Vision are specific to a given dataset, it would be inappropriate to compare scores across distinct datasets.

Here, we used these signatures to assess maturation state differences in recently identified murine white adipocyte precursors ASC1 and ASC2 (Rondini and Granneman 2020; Deutsch et al. 2020). ASC1 and ASC2 were recently identified as two distinct preadipocyte-types in multiple mouse scRNA-seq studies (Fig. 4A to 4C), with *in vitro* studies reflecting differential adipogenic capacity, and hence differential therapeutic capacity, within these cell populations ((Merrick et al. 2019; Schwalie et al. 2018; Hepler et al. 2018). However, *in vivo* studies revealed a transition of ASC2 into ASC1 prior to becoming adipocytes (Merrick et al. 2019), thereby suggesting a less committed progenitor state for ASC2 cells as compared to ASC1. Overall, it remains unclear whether ASC1 and ASC2 cells are distinct cell-types or cells with different adipogenic maturation states. Hence, we used our gene signatures to determine whether maturation state differed between these cells. We first validated recovery of existing maturation state differences between preadipocytes and mature adipocytes using primary adipose tissue snRNA-seq datasets (Sun et al. 2020). As characterized in the original study, the snRNA-seq dataset revealed a primary cluster of preadipocytes and adipocytes each in WAT and BAT, as marked by the expression of marker genes *DCLK1* and *ADIPOQ* (Fig. S4A and S4B). Gene signatures in WAT and BAT revealed a significantly higher score for mature white and brown adipocytes, as compared to respective preadipocytes (Fig. S4C and S4D), thereby validating the applicability of our signatures to identify maturation state differences. Next, we applied our signatures to three distinct scRNA-seq studies that identified ASC1 and ASC2 precursor cells (Fig. 4A to 4C) in murine WAT (Merrick et al. 2019; Schwalie et al. 2018; Hepler et al. 2018). In all three studies, the signature score was significantly higher in ASC1 cells compared to ASC2 (Fig. 4D to 4F), further corroborating the hypothesis that the two precursors are cells at different stages of adipogenic maturation, rather than being two distinct cell-types.

**Fig. 4.**
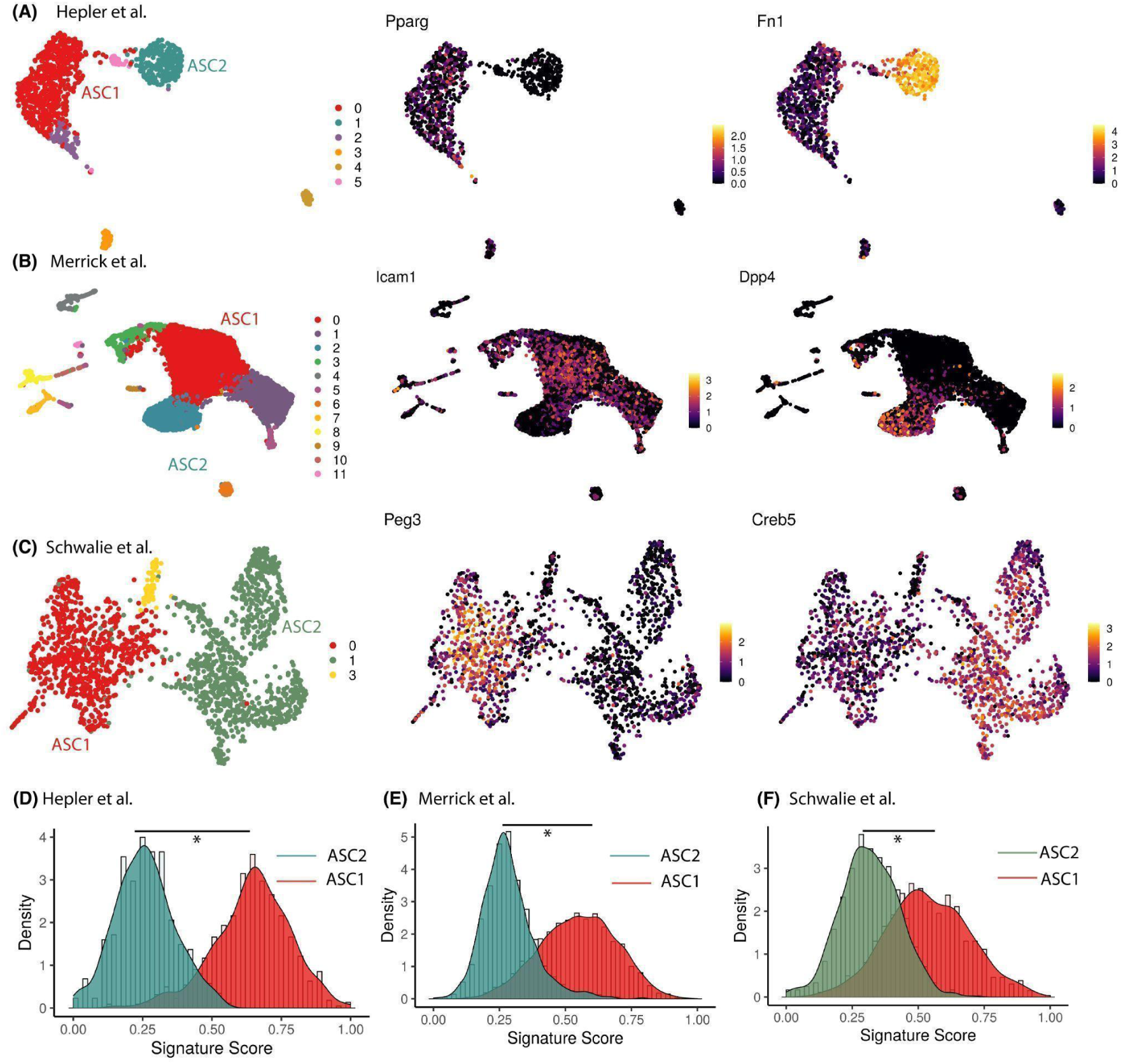
Cell maturation state assessment for murine white adipocyte precursors ASC1 and ASC2 (A) to (C) UMAP visualization of murine WAT scRNA-seq datasets from different studies colored by cluster (left panel), ASC1 marker expression (middle panel) and ASC2 marker expression (right panel) **(D) to (F)** Distribution of gene signature score for ASC2 and ASC1 cells from different studies. Score was scaled to vary from 0 to 1.

#### Differential expression and signature score analyses reveal inhibited adipogenic expansion in obese metabolic phenotype

In healthy adipogenic expansion, new adipocytes are derived from *de novo* differentiation of adipose progenitors or stem cells (hyperplastic expansion). In obesity, however, dysregulated adipogenesis results in excess fat being stored in existing adipocytes, resulting in their enlarged and inflamed phenotype (hypertrophic expansion; (Jo et al. 2009). Here, using our dataset, we investigated which of the genes dynamically regulated during healthy adipogenesis are differentially expressed in lean vs obese individuals.

First, we worked with 3 publicly available WAT datasets, 2 of which profiled gene expression using bulk RNA-sequencing (Zhou et al. 2020; Fisk et al. 2021) and the third using microarray (Arner et al. 2012). We observed that many dynamically regulated genes in our study were differentially expressed between obese and lean samples (Table S4). Particularly, a majority of Module 1 and Module 2 genes were differentially enriched in obese samples in all 3 studies (Fig. 5A and Fig. S5A). This is expected since downregulated genes primarily include CAMs and fibrillar ECM components, which exhibit increased production and accumulation during adipose tissue fibrosis in obesity (DeBari and Abbott 2020). Conversely, Module 4 and Module 5 genes, which primarily included adipogenic and lipogenic markers, were preferentially differentially enriched in lean samples in all 3 studies (Fig. 5A and Fig. S5A). This observation was consistent with previous reports demonstrating downregulation of adipogenic genes in obesity (Dubois et al. 2006; Nadler et al. 2000), possibly due to hypertrophic adipose tissue expansion, as well as adipocyte dedifferentiation (Song and Kuang 2019).

**Fig. 5.**
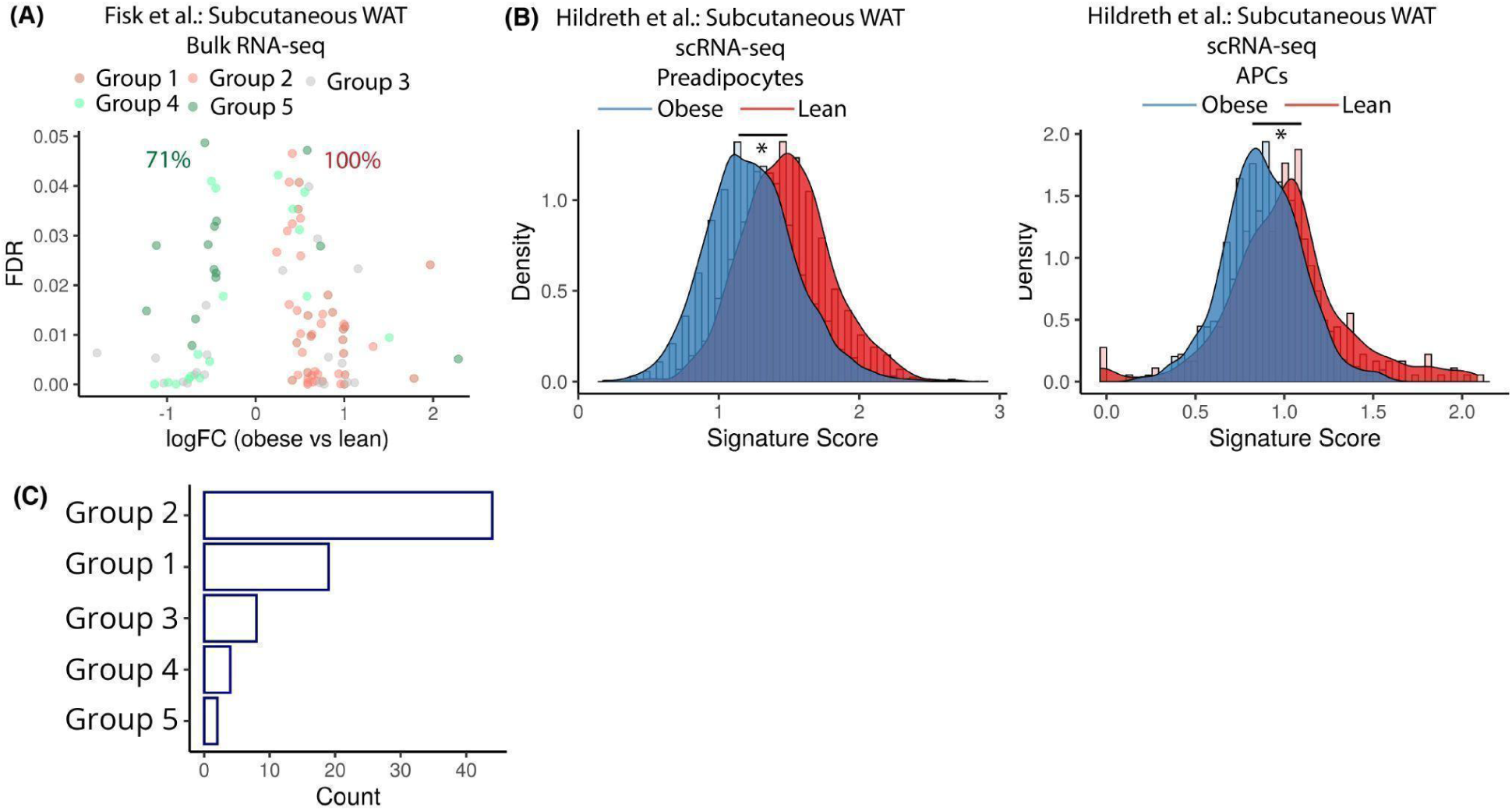
Implication of dynamically regulated genes during human adipogenesis in obesity. **(A)** Volcano plot of dynamically regulated genes DE between lean vs obese bulk RNA-seq samples. Each dot is a gene colored by its original gene module annotation. The number in green indicates percent of total DE genes in Module 4 and 5 that are enriched in lean samples. The number in red indicates percent of total DE genes in Module 1 and 2 that are enriched in obese samples. **(B)** Distribution of adipogenic gene signature scores between lean and obese patient in Preadipocytes (left panel) and Adipocyte precursors (APCs; right panel). **(C)** Distribution of all dynamically regulated genes during white adipogenesis that are also reported in GWAS datasets according to their annotation using gene modules from the current study.

At the bulk level, our results suggested there may be adipogenic inhibition phenotypes in samples from individuals with obesity. However, it remains unclear whether similar molecular differences also exist at the preadipocyte level, since cell-type level resolution is lost in bulk measurements. Using a recently published scRNA-seq WAT dataset with comparison of lean vs obese humans (Hildreth et al. 2021), we utilized our white lineage gene signature to investigate differences in maturation state at the preadipocyte level. The original study identified two distinct clusters, one of adipocyte precursors (APCs), and another of preadipocytes (Fig. S5B). Preadipocytes are known to be derived from APCs as part of the differentiation process, and this trend was also recovered by our signature analysis (Fig. S5C) Notably, the original study reported a lower fraction of APCs, and a higher fraction of preadipocytes in lean patients, suggesting an accelerated development from APCs to preadipocytes in lean individuals. Our signature analysis revealed a significantly higher maturation score for both preadipocytes and APCs in lean samples (Fig. 5B), suggesting existence of adipogenic inhibition amongst the two cell populations associated with obesity.

Besides gene expression profiling, genome-wide association studies have also been critical in linking genetic variants to obesity and metabolic disease risk. Here, we took an integrative approach to further analyze our adipogenic-molecular findings in light of recent GWAS studies and asked whether genes dynamically regulated in human adipogenesis are also linked to metabolic traits associated with increased genetic risk for obesity.

Using publicly available GWAS datasets (see Methods), we identified SNPs that are associated with metabolic traits such as BMI, waist circumference, and hip circumference (Locke et al. 2015; Shungin et al. 2015). In total, we analyzed datasets from 19 studies identifying over 1000 SNPs localized within the genic regions of 984 distinct genes. 77 of those 984 genes were temporally regulated during differentiation of white preadipocytes in our dataset. 44 of the 77 genes belonged to Module-2 which was associated with ECM and cytoskeletal remodeling during adipogenesis (Fig. 5C). This observation agrees with dysfunctional ECM remodeling being a hallmark of obesity, in which excessive lipid accumulation in adipocytes provokes an excess of deposition of ECM components such as collagens, elastin, and fibronectin in adipose tissue (Lin, Chun, and Kang 2016; Ruiz-Ojeda et al. 2019). Of the 77 genes, 15 genes were transcription factors that were both temporally regulated during white adipogenesis and associated with metabolic disease risk traits (Fig. S5D). The transcription factor *EBF1*, which was exclusively regulated during brown adipogenesis, was also associated with metabolic disease risk traits. This observation provides further evidence for a potentially thermogenic role of EBF1 (Angueira et al. 2020).

## DISCUSSION

In this study, we decipher the transcriptional dynamics of human white and brown fat development using an *in vitro* model system. Utilization of the *in vitro* model system enables high-resolution sampling of the entirety of adipogenic cell maturation states. We ordered differentiating adipocytes from the white and brown lineages by their progression through differentiation based on an adipogenic gene signature score. After cellular ordering, unsupervised gene clustering revealed five transcriptional modules with distinct expression dynamics that result in the generation of mature white and brown adipocytes. Through trajectory comparison, we identified novel TFs potentially involved in regulation of adipogenic or thermogenic transcriptional responses.

Integrative analysis of differentiating brown preadipocytes identified a precursor population with non-adipogenic response (Preadipocyte-1). Previously, non-adipogenic cell-types called Aregs were identified, which inhibit adipogenesis of other APCs in a paracrine fashion (Schwalie et al. 2018). However, Aregs are CD142+ which, in our dataset, was not expressed in the Preadipocyte-1 population. Recently, multiple scRNA-seq studies reported the existence of two primary APC populations in mice referred to as Asc1 and Asc2 (Deutsch et al. 2020; Rondini and Granneman 2020). Like adipogenic brown preadipocytes (Preadipocyte-2), Asc2 exhibited pro-inflammatory and pro-fibrotic phenotype and positive expression of genes *PI16* and *MFAP5*. Functional investigations into Asc2 and Asc1 cells revealed that Asc2 inhibited the differentiation of Asc1 cells *in vitro* (Rondini and Granneman 2020). Therefore, it is plausible that Preadipocyte-2 cells identified in our study may be functioning in a manner like Asc2, possibly to maintain adipocyte turnover. However, further functional investigations are necessary to validate the existence and functionality of multiple progenitor cell types in primary brown adipose tissue.

Within both white and brown adipogenesis, the early underlying transcriptional program was revealed to be centered around ECM remodeling. Although down-regulation of fibrillar ECM and up-regulation of basement membrane ECM has been demonstrated during 3T3-L1 adipogenesis, the dynamics of ECM reorganization and its regulation during human adipogenesis are not well understood. Our findings provide a high-resolution view into the expression dynamics of specific ECM components during healthy adipose tissue expansion, as well as potential ECM remodeling regulated by an interplay of metalloproteases and its inhibitors. In future investigations, our dataset could be used as a reference for analyses into differential expression dynamics of ECM components or metalloproteases in healthy vs metabolically diseased states.

An important aspect of our model system is the isolation of white and brown preadipocytes from a single individual, and a single anatomical location. Such a paired isolation strategy eliminates the effect of genetic variation, and enables investigation not just within, but also across white and brown lineages. Using our model system, we identified *RFX8* and *SOX5* as new TFs with potential adipogenic regulatory activity in both white and brown fat development. Moreover, based on exclusive regulation/enrichment in brown adipogenesis, we also identified novel TFs *ZNF117* and *IRX6* with potential involvement in thermogenic regulation. Future functional investigations would be required to confirm the regulatory roles of these TFs in healthy as well as diseased conditions. Ultimately, identification of novel TFs helps discern specific pharmacological targets for stimulating metabolically healthy, as well as thermogenic fat development.

In order to assign a pseudo-time to differentiating human preadipocytes, we identified gene sets which defined transcriptional signatures specifically associated with white or brown fat development. Scores for these transcriptional signatures could be utilized as a quantitative metric to investigate differences in cellular maturity across varying metabolic conditions, and for different anatomical locations. Such an analysis could help us better understand the differential roles of fat depots towards obesity development and progression. Moreover, our gene signature provides a targeted list of temporally, and biologically relevant genes, which could be specifically profiled using techniques such as spatial transcriptomics, or *in situ* hybridization/sequencing (Burgess 2019; Asp, Bergenstråhle, and Lundeberg 2020), to better understand the spatial organization of adipose tissue development in primary samples. Finally, our gene signature could also be used to understand adipocyte dedifferentiation, and investigate molecular differences between dedifferentiated preadipocytes, and differentiating preadipocytes, at similar stages of maturity.

Our study design also informs on the future directions for comprehensively investigating molecular regulation of human adipogenesis. In this work, adipogenic transcriptional dynamics were investigated using an immortalized, *in vitro* system of human white and brown preadipocytes. This dataset will serve as reference to further understand molecular circuitry of fat development in primary adipose tissue samples. Additionally, the currently employed *in vitro* model system was isolated from a single individual, and from a single anatomical location. And as such, the transcriptional landscape generated here could be made even more comprehensive by isolating similar model systems across multiple individuals and depot locations. During QC of snRNA-seq libraries analyzed in this investigation, we observed varying levels of background mRNA contamination in individual samples. Single-nuclei extraction involves breaking apart the cellular matrix to isolate nuclei, which releases high amounts of debris and cytoplasmic mRNA. During droplet-based single nuclei isolation, this debris gets encapsulated in the droplet along with the nuclei, leading to background mRNA contamination (Alvarez et al. 2020; Fleming, Marioni, and Babadi 2019). This varying mRNA contamination makes it challenging to identify nuclei that are at similar stages of differentiation but distributed across different days of harvest (different single-nuclei libraries). scRNA-seq dataset integration algorithms do mitigate this challenge partially. However, development of integration strategies taking into account varying background mRNA distribution in snRNA-seq datasets would provide more accurate insights into the gene expression dynamics during differentiation processes.

Overall, our study takes the first steps towards understanding the nature of adipogenic differentiation at a high temporal and cellular resolution in humans. These findings will therefore serve as a resource for multiple efforts into investigating adipose tissue biology in health, as well as disease, ultimately enabling newer therapeutics for improved clinical tackling of a variety of metabolic disorders. Fundamentally, our dataset provides a high-resolution resource for mapping different adipogenic cell-types and states in the human adipose tissue and therefore, serves as a reference for undertakings such as mapping the Human Cell Atlas (Regev et al. 2017).

## METHODS

### Preadipocyte culture and adipogenic differentiation

Detailed protocol for maintenance, cryopreservation, and differentiation of white and brown preadipocytes are outlined in a different study (Shamsi and Tseng 2017). Briefly, for culturing preadipocytes, cells were grown in DMEM medium (Corning, 10-017-CV) supplemented with 10% vol/vol FBS and containing 1% vol/vol Penicillin-Streptomycin (Gibco). Cell culture was maintained at 37°C in a humidified incubator containing 5% vol/vol CO2. 80% confluent cells were passaged using 0.25% trypsin with 0.1% EDTA (Gibco, 25200-056) for a 1:3 split in a new 100 mm cell culture dish (Corning).

Prior to adipogenic differentiation, white preadipocytes were allowed to grow up to 100% confluence in a 100 mm cell culture dish (Corning). After 48 hours at 100% confluence, growth media was replaced with adipogenic induction media every 48 hours for the next 20 days. Induction media was prepared by adding 1 mL FBS, 500 μl Penicillin-Streptomycin, 15 μl human Insulin (0.5 μM, Sigma-Aldrich, I2643-50MG), 10 μl T3 (2 nM, Sigma-Aldrich,T6397-100MG), 50 μl Biotin (33 μM, Sigma-Aldrich, B4639-100MG), 100 μl Pantothenate (17 μM, Sigma-Aldrich, P5155-100G), 1 μl Dexamethasone (0.1 μM, Sigma-Aldrich, D2915-100MG), 500 μl IBMX (500 μM, Sigma-Aldrich, I7018-100mg), and 12.5 μl Indomethacin (30 μM, Sigma-Aldrich, I7378-5G) to 48.5 mL DMEM medium and sterile filter.

### Nuclei isolation from differentiating preadipocytes and snRNA-seq

Nuclei were isolated from differentiating white and brown preadipocytes using an NP-40 based lysis buffer: To 14.7 mL nuclease-free water (Qiagen), 150 uL of Tris-Hydrochloride (Sigma, T2194), 30 uL of Sodium Chloride (5M; Sigma, 59222C), 45 uL of Magnesium Chloride (1M; Sigma, M1028), and 75 uL of NP-40 (Sigma, 74385) was added. Nuclei harvested on day-0 were isolated from preadipocytes prior to adipogenic differentiation. Two 100 mm dishes were used for nuclei isolation from each preadipocyte type. 500 uL of NP-40 based lysis buffer was added to each 100 mm dish and a cell scraper was employed to release adherent cells from the plates. On day 10, 15, and 20, where cells had visible lipid droplet accumulation, dounce homogenizer was used on scraped out cells to separate the lipids. Cells were then incubated with the lysis buffer for 5 minutes on ice in a pre-chilled 15 mL falcon tube. Cells were washed with ice-cold PBS supplemented with .2 U/uL RNase Inhibitor (Protector RNase Inhibitor; henceforth called wash buffer) 4 times by centrifuging at 500 rcf for 5 minutes at 4C. Wash buffer was aspirated after the final round of centrifugation and nuclei were resuspended in the ice-cold wash buffer and filtered using a 40 um cell strainer. Final concentration was adjusted to ∼1000 nuclei/uL using a hemocytometer for downstream sequencing. Nuclei were also stained using 0.08% trypan blue dye to assess nuclear membrane integrity under brightfield imaging. For nuclear isolation on day 10, 15, and 20, the same protocol was implemented as mentioned above with the modification of using 1 mL lysis buffer for each 100 mm dish.

After preparing nuclei suspension, isolation was performed on the 10x Chromium platform and libraries prepared as per the manufacturer’s protocol using v3 sequencing chemistry. All final libraries were sequenced on the Illumina NovaSeq platform to the following specifications:

### Sequencing data QC and analysis

In total, we had 5 libraries each for the white and brown adipogenesis dataset. For each library, empty droplets were removed using CellBender (Fleming et al., 2019), and doublets were removed using Scrublet (Wolock, Lopez, and Klein 2019) or DoubletDetection (Gayoso et al. 2019). Using Seurat, low-quality clusters such as clusters with high MT content, clusters with cellular debris (as marked by the enrichment of translation terms in GO analysis), clusters enriched for empty/doublet barcodes were removed from downstream analysis. Integration of all 5 time points for white and brown dataset was performed using scVI-tools (Gayoso et al., 2021). Post-integration, Seurat was used for unsupervised clustering, and differential gene expression analysis. Transcription factor enrichment analysis was performed using ChEA3.

### Pseudo-temporal Ordering

#### Slingshot Analysis

For white and brown adipogenesis dataset, the integrated Seurat object was clustered at resolution = 0.4. Slingshot was then used to infer the trajectory using the cluster with the highest contribution from day 0 as the starting cluster. For identifying temporally regulated genes, cells were clustered into 6 equally spaced pseudo-temporal bins. DGE was then performed for each bin against the 1st and last bin, and all genes with logFC $>$ 1 and FDR $<$ 0.05 were considered as temporally regulated. For identifying monotonically increasing genes when cells are ordered using Slingshot, genes were clustered using the ComplexHeatmap package, with k-means clustering algorithm, and the number of clusters set to 5.

#### Vision analysis

For both white and brown adipogenesis dataset, Vision was used to assign a score to each cell in the integrated Seurat object using the “Hallmark_Adipogenesis” MSigDB signature. This score was used as a proxy for pseudotime. For identifying temporally regulated genes in white adipogenesis dataset, cells were distributed into bins defined using the command *cut_points=c(-Inf,seq(0.15,0.6,0.15),Inf)*. For the brown adipogenesis dataset, cells were distributed into bins defined using the command *cut_points=c(-Inf,0.2,0.3,seq(0.4,0.6,0.2),Inf)*. Temporally regulated genes were identified using the same strategy as defined above. For identifying monotonically increasing genes when cells are ordered using Vision, genes were clustered using the ComplexHeatmap package, with k-means clustering algorithm, and the number of clusters set to 5.

#### Identifying lineage-specific gene signatures

Once monotonically increasing genes were identified using both Vision and Slingshot, the intersection of the two was taken to define lineage-specific gene signatures. These signatures were used as input to Vision to assign a score to differentiating white and brown preadipocytes, and used as a proxy for pseudotime.

#### Gene Module Clustering

For identifying temporally regulated genes in white dataset, when cells are ordered using lineage-specific gene signatures, cells were distributed into bins defined using the command *cut_points=c(-Inf,0.1,seq(0.3,0.8,0.1),Inf*. DGE was then performed for each bin against the 1st and last bin, and all genes with logFC $>$ 1 and FDR $<$ 0.05 were considered as temporally regulated. Clustering for genes was performed using Seurat with resolution set to 0.5. For the brown adipogenesis dataset, the same steps were used with *cut_points* defined using the command *cut_points=c(-Inf,0.15,0.225,seq(0.3,0.7,0.2),Inf)*.

### GWAS Analysis

The GWAS dataset was downloaded from the GWAS catalog (gwas_catalog_v1.0-associations_e100_r2021-04-20.tsv) and subset to metabolic traits defined in Locke et al. and Shungin et al. The catalog was further subset to SNPs that were mapped to a single gene.

### Bulk RNA-seq and Microarray Analysis

RNA-seq datasets were downloaded from the GEO Accession Viewer using Accession# GSE25401 and GSE162653. Microarray data was downloaded from the journal’s website (oby22950-sup-0007-TableS1.xlsx). Differential expression analysis for RNA-seq datasets was performed using the DESeq package in R.

### Adipogenic signature scoring analysis in publicly available scRNA-seq/snRNA-seq datasets

For assessing the maturation of cells in publicly available scRNA-seq datasets, Vision was used to assign score to cells using our lineage-specific signatures. Datasets were downloaded from accession numbers mentioned in the original manuscript.

## DATA ACCESS

Data related to this study is available upon request to the corresponding author.

## ACKNOWLEDGEMENTS

This publication was supported by the National Institute of General Medical Sciences of the National Institutes of Health under award number R35GM124916. This publication was also supported by grant number CZF2019-002454 from the Chan Zuckerberg Foundation and grant numbers R01DK102898 and K01DK125608 from the National Institutes of Health. This project has been made possible in part by grant numbers 2019-002454 and 2022-246198 from the Chan Zuckerberg Initiative DAF, an advised fund of Silicon Valley Community Foundation. A.S. is a Chan-Zuckerberg Biohub Investigator and a Pew Scholar in the Biomedical Sciences, supported by the Pew Charitable Trusts. A.G. is supported by the University of California at Berkeley Lloyd Fellowship in Bioengineering.

## REFERENCES

Ahmadian, Maryam, Marcia J. Abbott, Tianyi Tang, Carolyn S. S. Hudak, Yangha Kim, Matthew Bruss, Marc K. Hellerstein, et al. 2011. “Desnutrin/ATGL Is Regulated by AMPK and Is Required for a Brown Adipose Phenotype.” Cell Metabolism 13 (6): 739–48.

Al Hasan, Mohammad, Patricia E. Martin, Xinhua Shu, Steven Patterson, and Chris Bartholomew. 2021. “Type III Collagen Is Required for Adipogenesis and Actin Stress Fibre Formation in 3T3-L1 Preadipocytes.” Biomolecules 11 (2): 1–17.

Alvarez, Marcus, Elior Rahmani, Brandon Jew, Kristina M. Garske, Zong Miao, Jihane N. Benhammou, Chun Jimmie Ye, et al. 2020. “Enhancing Droplet-Based Single-Nucleus RNA-Seq Resolution Using the Semi-Supervised Machine Learning Classifier DIEM.” Scientific Reports 10 (1): 1–16.

Ambele, Melvin Anyasi, Carla Dessels, Chrisna Durandt, and Michael Sean Pepper. 2016. “Genome-Wide Analysis of Gene Expression during Adipogenesis in Human Adipose-Derived Stromal Cells Reveals Novel Patterns of Gene Expression during Adipocyte Differentiation.” Stem Cell Research 16 (3): 725–34.

Angueira, Anthony R., Suzanne N. Shapira, Jeff Ishibashi, Samay Sampat, Jaimarie Sostre-Colón, Matthew J. Emmett, Paul M. Titchenell, Mitchell A. Lazar, Hee-Woong Lim, and Patrick Seale. 2020. “Early B Cell Factor Activity Controls Developmental and Adaptive Thermogenic Gene Programming in Adipocytes.” Cell Reports 30 (9): 2869–78.e4.

Arner, Erik, Niklas Mejhert, Agné Kulyté, Piotr J. Balwierz, Mikhail Pachkov, Mireille Cormont, Silvia Lorente-Cebrián, et al. 2012. “Adipose Tissue microRNAs as Regulators of CCL2 Production in Human Obesity.” Diabetes 61 (8): 1986–93.

Asp, Michaela, Joseph Bergenstråhle, and Joakim Lundeberg. 2020. “Spatially Resolved Transcriptomes-next Generation Tools for Tissue Exploration.” BioEssays: News and Reviews in Molecular, Cellular and Developmental Biology 42 (10): e1900221.

Betz, Matthias J., and Sven Enerbäck. 2015. “Human Brown Adipose Tissue: What We Have Learned So Far.” Diabetes 64 (7): 2352–60.

Bi, Pengpeng, Tizhong Shan, Weiyi Liu, Feng Yue, Xin Yang, Xin Rong Liang, Jinghua Wang, et al. 2014. “Inhibition of Notch Signaling Promotes Browning of White Adipose Tissue and Ameliorates Obesity.” Nature Medicine 20 (8): 911–18.

Birnbaum, Kenneth D. 2018. “Power in Numbers: Single-Cell RNA-Seq Strategies to Dissect Complex Tissues.” Annual Review of Genetics. https://doi.org/10.1146/annurev-genet-120417-031247.

Blondin, Denis P., Soren Nielsen, Eline N. Kuipers, Mai C. Severinsen, Verena H. Jensen, Stéphanie Miard, Naja Z. Jespersen, et al. 2020. “Human Brown Adipocyte Thermogenesis Is Driven by β2-AR Stimulation.” Cell Metabolism 32 (2): 287–300.e7.

Burgess, Darren J. 2019. “Spatial Transcriptomics Coming of Age.” Nature Reviews. Genetics.

Burl, Rayanne B., Vanesa D. Ramseyer, Elizabeth A. Rondini, Roger Pique-Regi, Yun Hee Lee, and James G. Granneman. 2018. “Deconstructing Adipogenesis Induced by β3-Adrenergic Receptor Activation with Single-Cell Expression Profiling.” Cell Metabolism 28 (2): 300–309.e4.

Caserta, F., T. Tchkonia, V. N. Civelek, M. Prentki, N. F. Brown, J. D. McGarry, R. A. Forse, B. E. Corkey, J. A. Hamilton, and J. L. Kirkland. 2001. “Fat Depot Origin Affects Fatty Acid Handling in Cultured Rat and Human Preadipocytes.” American Journal of Physiology. Endocrinology and Metabolism 280 (2): E238–47.

Cederberg, Anna, Line M. Gronning, B. Ahrén, Kjetil Taskén, Peter Carlsson, and Sven Enerbäck. 2001. “FOXC2 Is a Winged Helix Gene That Counteracts Obesity, Hypertriglyceridemia, and Diet-Induced Insulin Resistance.” Cell 106 (5): 563–73.

Cero, Cheryl, Hannah J. Lea, Kenneth Y. Zhu, Farnaz Shamsi, Yu-Hua Tseng, and Aaron M. Cypess. 2021. “β3-Adrenergic Receptors Regulate Human Brown/beige Adipocyte Lipolysis and Thermogenesis.” JCI Insight 6 (11). https://doi.org/10.1172/jci.insight.139160.

Chen, Junye, Yi Lu, Mengyuan Tian, and Qiren Huang. 2019. “Molecular Mechanisms of foxo1 in Adipocyte Differentiation.” Journal of Molecular Endocrinology. https://doi.org/10.1530/JME-18-0178.

Choy, Lisa, Jeremy Skillington, and Rik Derynck. 2000. “Roles of Autocrine TGF-β Receptor and Smad Signaling in Adipocyte Differentiation.” The Journal of Cell Biology 149 (3): 667–81.

Côté Julie Anne, Julie Lessard, Mélissa Pelletier, Simon Marceau, Odette Lescelleur, Julie Fradette, and André Tchernof. 2017. “Role of the TGF-β Pathway in Dedifferentiation of Human Mature Adipocytes.” FEBS Open Bio 7 (8): 1092–1101.

Cypess, Aaron M., Andrew P. White, Cecile Vernochet, Tim J. Schulz, Ruidan Xue, Christina A. Sass, Tian Liang Huang, et al. 2013. “Anatomical Localization, Gene Expression Profiling and Functional Characterization of Adult Human Neck Brown Fat.” Nature Medicine 19 (5): 635–39.

DeBari, Megan K., and Rosalyn D. Abbott. 2020. “Adipose Tissue Fibrosis: Mechanisms, Models, and Importance.” International Journal of Molecular Sciences 21 (17). https://doi.org/10.3390/ijms21176030.

DeTomaso, David, Matthew G. Jones, Meena Subramaniam, Tal Ashuach, Chun J. Ye, and Nir Yosef. 2019. “Functional Interpretation of Single Cell Similarity Maps.” Nature Communications 10 (1). https://doi.org/10.1038/s41467-019-12235-0.

Deutsch, Alana, Daorong Feng, Jeffrey E. Pessin, and Kosaku Shinoda. 2020. “The Impact of Single-Cell Genomics on Adipose Tissue Research.” International Journal of Molecular Sciences. MDPI AG. https://doi.org/10.3390/ijms21134773.

Dubois, Severine G., Leonie K. Heilbronn, Steven R. Smith, Jeanine B. Albu, David E. Kelley, Eric Ravussin, and Look AHEAD Adipose Research Group. 2006. “Decreased Expression of Adipogenic Genes in Obese Subjects with Type 2 Diabetes.” Obesity 14 (9): 1543–52.

Ehrlund, Anna, Niklas Mejhert, Christel Björk, Robin Andersson, Agné Kulyté, Gaby Åström, Masayoshi Itoh, et al. 2017. “Transcriptional Dynamics during Human Adipogenesis and Its Link to Adipose Morphology and Distribution.” Diabetes 66 (1): 218–30.

Fan, Wu Qiang, Takeshi Imamura, Noriyuki Sonoda, Dorothy D. Sears, David Patsouris, Jane J. Kim, and Jerrold M. Olefsky. 2009. “FOXO1 Transrepresses Peroxisome Proliferator-Activated Receptor γ Transactivation, Coordinating an Insulin-Induced Feed-Forward Response in Adipocytes.” The Journal of Biological Chemistry 284 (18): 12188–97.

Fisk, Helena L., Caroline E. Childs, Elizabeth A. Miles, Robert Ayres, Paul S. Noakes, Carolina Paras-Chavez, Ondrej Kuda, et al. 2021. “Dysregulation of Endocannabinoid Concentrations in Human Subcutaneous Adipose Tissue in Obesity and Modulation by Omega-3 Polyunsaturated Fatty Acids.” Clinical Science 135 (1): 185–200.

Fleming, Stephen J., John C. Marioni, and Mehrtash Babadi. 2019. “CellBender Remove-Background: A Deep Generative Model for Unsupervised Removal of Background Noise from scRNA-Seq Datasets.” bioRxiv, Oober, 791699.

Fontaine, Coralie, Wendy Cousin, Magali Plaisant, Christian Dani, and Pascal Peraldi. 2008. “Hedgehog Signaling Alters Adipocyte Maturation of Human Mesenchymal Stem Cells.” Stem Cells 26 (4): 1037–46.

Gayoso, Adam, Romain Lopez, Galen Xing, Pierre Boyeau, Katherine Wu, Michael Jayasuriya, Edouard Melhman, et al. 2021. “Scvi-Tools: A Library for Deep Probabilistic Analysis of Single-Cell Omics Data.” bioRxiv. https://doi.org/10.1101/2021.04.28.441833.

Gayoso, Adam, Jonathan Shor, Ambrose J. Carr, Roshan Sharma, and Dana Pe’er. 2019. “JonathanShor/DoubletDetection: HOTFIX: Correct Setup.py Installation,” August. https://doi.org/10.5281/ZENODO.3376859.

Gerin, Isabelle, Guido T. Bommer, Martin E. Lidell, Anna Cederberg, Sven Enerback, and Ormond A. McDougald. 2009. “On the Role of FOX Transcription Factors in Adipocyte Differentiation and Insulin-Stimulated Glucose Uptake.” The Journal of Biological Chemistry 284 (16): 10755–63.

Gubelmann, Carine, Petra C. Schwalie, Sunil K. Raghav, Eva Röder, Tenagne Delessa, Elke Kiehlmann, Sebastian M. Waszak, et al. 2014. “Identification of the Transcription Factor ZEB1 as a Central Component of the Adipogenic Gene Regulatory Network.” eLife 3 (August 2014): 1–30.

Gupta, Anushka, Farnaz Shamsi, Nicolas Altemos, Gabriel F. Dorlhiac, Aaron M. Cypess, Andrew P. White, Mary Elizabeth Patti, Yu-Hua Tseng, and Aaron Streets. 2021. “Characterization of Transcript Enrichment and Detection Bias in Single-Nuclei RNA-Seq for Mapping of Distinct Human Adipocyte Lineages.” bioRxiv, March, 2021.03.24.435852.

Han, Xiaoping, Renying Wang, Yincong Zhou, Lijiang Fei, Huiyu Sun, Shujing Lai, Assieh Saadatpour, et al. 2018. “Mapping the Mouse Cell Atlas by Microwell-Seq.” Cell 172 (5): 1091–1107.e17.

Harms, Matthew, and Patrick Seale. 2013. “Brown and Beige Fat: Development, Function and Therapeutic Potential.” Nature Medicine 19 (10): 1252–63.

Heglind, Mikael, Anna Cederberg, Jorge Aquino, Guilherme Lucas, Patrik Ernfors, and Sven Enerbäck. 2005. “Lack of the Central Nervous System-and Neural Crest-Expressed Forkhead Gene Foxs1 Affects Motor Function and Body Weight.” Molecular and Cellular Biology 25 (13): 5616–25.

Hepler, Chelsea, Bo Shan, Qianbin Zhang, Gervaise H. Henry, Mengle Shao, Lavanya Vishvanath, Alexandra L. Ghaben, et al. 2018. “Identification of Functionally Distinct Fibro-Inflammatory and Adipogenic Stromal Subpopulations in Visceral Adipose Tissue of Adult Mice.” eLife 7 (September). https://doi.org/10.7554/eLife.39636.

Herzig, Sébastien, and Reuben J. Shaw. 2018. “AMPK: Guardian of Metabolism and Mitochondrial Homeostasis.” Nature Reviews Molecular Cell Biology. Nature Publishing Group. https://doi.org/10.1038/nrm.2017.95.

Hildreth, Andrew D., Feiyang Ma, Yung Yu Wong, Ryan Sun, Matteo Pellegrini, and Timothy E. O’Sullivan. 2021. “Single-Cell Sequencing of Human White Adipose Tissue Identifies New Cell States in Health and Obesity.” Nature Immunology 22 (5): 639–53.

Hrckulak, Dusan, Lucie Janeckova, Lucie Lanikova, Vitezslav Kriz, Monika Horazna, Olga Babosova, Martina Vojtechova, Katerina Galuskova, Eva Sloncova, and Vladimir Korinek. 2018. “Wnt Effector TCF4 Is Dispensable for Wnt Signaling in Human Cancer Cells.” Genes 9 (9). https://doi.org/10.3390/genes9090439.

Hurtado Del Pozo, Carmen Gregorio Vesperinas-García, Miguel Ángel Rubio, Ramón Corripio-Sánchez, Antonio J. Torres-García, Maria Jesus Obregon, and Rosa María Calvo. 2011. “ChREBP Expression in the Liver, Adipose Tissue and Differentiated Preadipocytes in Human Obesity.” Biochimica et Biophysica Acta - Molecular and Cell Biology of Lipids 1811 (12): 1194–1200.

Hussain, Mohammed Faiz, Anna Roesler, and Lawrence Kazak. 2020. “Regulation of Adipocyte Thermogenesis: Mechanisms Controlling Obesity.” The FEBS Journal 287 (16): 3370–85.

Ishihara, Yasuhiro, Mayumi Tsuji, and Christoph F. A. Vogel. 2018. “Suppressive Effects of Aryl-Hydrocarbon Receptor Repressor on Adipocyte Differentiation in 3T3-L1 Cells.” Archives of Biochemistry and Biophysics 642 (March): 75–80.

Jo, Junghyo, Oksana Gavrilova, Stephanie Pack, William Jou, Shawn Mullen, Anne E. Sumner, Samuel W. Cushman, and Vipul Periwal. 2009. “Hypertrophy And/or Hyperplasia: Dynamics of Adipose Tissue Growth.” Edited by Jason A. Papin. PLoS Computational Biology 5 (3): e1000324.

Karaman, Sinem, Maija Hollmén, Sun Young Yoon, H. Furkan Alkan, Kari Alitalo, Christian Wolfrum, and Michael Detmar. 2016. “Transgenic Overexpression of VEGF-C Induces Weight Gain and Insulin Resistance in Mice.” Scientific Reports 6 (August). https://doi.org/10.1038/srep31566.

Kim, Sooho, Chihoon Ahn, Naeun Bong, Senyon Choe, and Dong Kun Lee. 2015. “Biphasic Effects of FGF2 on Adipogenesis.” Edited by Marià Alemany. PloS One 10 (3): e0120073.

Kriszt, Rókus, Satoshi Arai, Hideki Itoh, Michelle H. Lee, Anna G. Goralczyk, Xiu Min Ang, Aaron M. Cypess, et al. 2017. “Optical Visualisation of Thermogenesis in Stimulated Single-Cell Brown Adipocytes.” Scientific Reports 7 (1). https://doi.org/10.1038/s41598-017-00291-9.

Lee, Haemi, Hyo Jung Kim, Yoo Jeong Lee, Min-Young Lee, Hyeonjin Choi, Hyemin Lee, and Jae-Woo Kim. 2012. “Krüppel-Like Factor KLF8 Plays a Critical Role in Adipocyte Differentiation.” PLoS ONE. https://doi.org/10.1371/journal.pone.0052474.

Li, Meihang, Zhenjiang Liu, Zhenzhen Zhang, Guannv Liu, Shiduo Sun, and Chao Sun. 2015. “miR-103 Promotes 3T3-L1 Cell Adipogenesis through AKT/mTOR Signal Pathway with Its Target Being MEF2D.” Biological Chemistry 396 (3): 235–44.

Lin, De, Tae-Hwa Chun, and Li Kang. 2016. “Adipose Extracellular Matrix Remodelling in Obesity and Insulin Resistance.” Biochemical Pharmacology 119 (November): 8–16.

Liu, Xuejiao, Christopher Cervantes, and Feng Liu. 2017. “Common and Distinct Regulation of Human and Mouse Brown and Beige Adipose Tissues: A Promising Therapeutic Target for Obesity.” Protein & Cell 8 (6): 446–54.

Locke, Adam E., Bratati Kahali, Sonja I. Berndt, Anne E. Justice, Tune H. Pers, Felix R. Day, Corey Powell, et al. 2015. “Genetic Studies of Body Mass Index Yield New Insights for Obesity Biology.” Nature 518 (7538): 197–206.

MacDougald, O. A., P. Cornelius, R. Liu, and M. D. Lane. 1995. “Insulin Regulates Transcription of the CCAAT/enhancer Binding Protein (C/EBP) α, β, and δ Genes in Fully-Differentiated 3T3-L1 Adipocytes.” The Journal of Biological Chemistry 270 (2): 647–54.

Mariman, E. C. M., and Ping Wang. 2010. “Adipocyte Extracellular Matrix Composition, Dynamics and Role in Obesity.” Cellular and Molecular Life Sciences. Springer. https://doi.org/10.1007/s00018-010-0263-4.

Ma, Xiuquan, Paul Lee, Donald J. Chisholm, and David E. James. 2015. “Control of Adipocyte Differentiation in Different Fat Depots; Implications for Pathophysiology or Therapy.” Frontiers in Endocrinology 6 (January): 1.

Medina-Gomez, Gema, Sarah Gray, and Antonio Vidal-Puig. 2007. “Adipogenesis and Lipotoxicity: Role of Peroxisome Proliferator-Activated Receptor γ (PPARγ) and PPARγcoactivator-1 (PGC1).” Public Health Nutrition 10 (10A): 1132–37.

Meex, Ruth C. R., Patrick Schrauwen, and Matthijs K. C. Hesselink. 2009. “Modulation of Myocellular Fat Stores: Lipid Droplet Dynamics in Health and Disease.” American Journal of Physiology. Regulatory, Integrative and Comparative Physiology 297 (4): R913–24.

Merrick, David, Alexander Sakers, Zhazira Irgebay, Chihiro Okada, Catherine Calvert, Michael P. Morley, Ivona Percec, and Patrick Seale. 2019. “Identification of a Mesenchymal Progenitor Cell Hierarchy in Adipose Tissue.” Science 364 (6438): eaav2501.

Morandi, E. M., R. Verstappen, M. E. Zwierzina, S. Geley, G. Pierer, and C. Ploner. 2016. “ITGAV and ITGA5 Diversely Regulate Proliferation and Adipogenic Differentiation of Human Adipose Derived Stem Cells.” Scientific Reports 6 (1): 1–14.

Mor-Yossef Moldovan Lisa, Maayan Lustig, Alex Naftaly, Mariya Mardamshina, Tamar Geiger, Amit Gefen, and Dafna Benayahu. 2019. “Cell Shape Alteration during Adipogenesis Is Associated with Coordinated Matrix Cues.” Journal of Cellular Physiology 234 (4): 3850–63.

Mota de Sá, Paula, Allison J. Richard, Hardy Hang, and Jacqueline M. Stephens. 2017. “Transcriptional Regulation of Adipogenesis.” Comprehensive Physiology 7 (2): 635–74.

Nadler, S. T., J. P. Stoehr, K. L. Schueler, G. Tanimoto, B. S. Yandell, and A. D. Attie. 2000. “The Expression of Adipogenic Genes Is Decreased in Obesity and Diabetes Mellitus.” Proceedings of the National Academy of Sciences of the United States of America 97 (21): 11371–76.

Nakajima, Ikuyo, Susumu Muroya, Ryo Ichi Tanabe, and Koichi Chikuni. 2002. “Extracellular Matrix Development during Differentiation into Adipocytes with a Unique Increase in Type V and VI Collagen.” Biology of the Cell / under the Auspices of the European Cell Biology Organization 94 (3): 197–203.

Regev, Aviv, Sarah A. Teichmann, Eric S. Lander, Ido Amit, Christophe Benoist, Ewan Birney, Bernd Bodenmiller, et al. 2017. “The Human Cell Atlas.” eLife 6 (December). https://doi.org/10.7554/eLife.27041.

Reusch, J. E., L. A. Colton, and D. J. Klemm. 2000. “CREB Activation Induces Adipogenesis in 3T3-L1 Cells.” Molecular and Cellular Biology 20 (3): 1008–20.

Rondini, Elizabeth A., and James G. Granneman. 2020. “Single Cell Approaches to Address Adipose Tissue Stromal Cell Heterogeneity.” The Biochemical Journal 477 (3): 583–600.

Rosen, Evan D., and Bruce M. Spiegelman. 2014. “Review What We Talk About When We Talk About Fat.” Cell 156 (1-2): 20–44.

Ross, Sarah E., Robin L. Erickson, Isabelle Gerin, Paul M. DeRose, Laszlo Bajnok, Kenneth A. Longo, David E. Misek, et al. 2002. “Microarray Analyses during Adipogenesis: Understanding the Effects of Wnt Signaling on Adipogenesis and the Roles of Liver X Receptor α in Adipocyte Metabolism.” Molecular and Cellular Biology 22 (16): 5989–99.

Ruiz-Ojeda, Francisco Javier, Andrea Méndez-Gutiérrez, Concepción María Aguilera, and Julio Plaza-Díaz. 2019. “Extracellular Matrix Remodeling of Adipose Tissue in Obesity and Metabolic Diseases.” International Journal of Molecular Sciences 20 (19). https://doi.org/10.3390/ijms20194888.

Sanchez-Gurmaches, Joan, Yuefeng Tang, Naja Zenius Jespersen, Martina Wallace, Camila Martinez Calejman, Sharvari Gujja, Huawei Li, et al. 2018. “Brown Fat AKT2 Is a Cold-Induced Kinase That Stimulates ChREBP-Mediated De Novo Lipogenesis to Optimize Fuel Storage and Thermogenesis.” Cell Metabolism 27 (1): 195–209.e6.

Sárvári Anitta Kinga, Elvira Laila Van Hauwaert, Lasse Kruse Markussen, Ellen Gammelmark, Ann Britt Marcher, Morten Frendø Ebbesen, Ronni Nielsen, Jonathan Richard Brewer, Jesper Grud Skat Madsen, and Susanne Mandrup. 2021. “Plasticity of Epididymal Adipose Tissue in Response to Diet-Induced Obesity at Single-Nucleus Resolution.” Cell Metabolism 33 (2): 437–53.e5.

Satish, Latha, J. Michael Krill-Burger, Phillip H. Gallo, Shelley Des Etages, Fang Liu, Brian J. Philips, Sudheer Ravuri, et al. 2015. “Expression Analysis of Human Adipose-Derived Stem Cells during in Vitro Differentiation to an Adipocyte Lineage.” BMC Medical Genomics 2015 8:1 8 (1): 1–12.

Schultz, Joshua R., Hua Tu, Alvin Luk, Joyce J. Repa, Julio C. Medina, Leping Li, Susan Schwendner, et al. 2000. “Role of LXRs in Control of Lipogenesis.” Genes and Development 14 (22): 2831–38.

Schwalie, Petra C., Hua Dong, Magda Zachara, Julie Russeil, Daniel Alpern, Nassila Akchiche, Christian Caprara, et al. 2018. “A Stromal Cell Population That Inhibits Adipogenesis in Mammalian Fat Depots.” Nature 559 (7712): 103–8.

Seale, P. 2010. “Transcriptional Control of Brown Adipocyte Development and Thermogenesis.” International Journal of Obesity 34 Suppl 1 (October): S17–22.

Seale, Patrick, Bryan Bjork, Wenli Yang, Shingo Kajimura, Sherry Chin, Shihuan Kuang, Anthony Scimè, et al. 2008. “PRDM16 Controls a Brown Fat/skeletal Muscle Switch.” Nature 454 (7207): 961–67.

Shan, Tizhong, Jiaqi Liu, Weiche Wu, Ziye Xu, and Yizhen Wang. 2017. “Roles of Notch Signaling in Adipocyte Progenitor Cells and Mature Adipocytes.” Journal of Cellular Physiology. Wiley-Liss Inc. https://doi.org/10.1002/jcp.25697.

Shao, Wei, and Peter J. Espenshade. 2012. “Expanding Roles for SREBP in Metabolism.” Cell Metabolism. NIH Public Access. https://doi.org/10.1016/j.cmet.2012.09.002.

Shapira, Suzanne N., and Patrick Seale. 2019. “Transcriptional Control of Brown and Beige Fat Development and Function.” Obesity 27 (1): 13–21.

Sharp, Louis Z., Kosaku Shinoda, Haruya Ohno, David W. Scheel, Emi Tomoda, Lauren Ruiz, Houchun Hu, et al. 2012. “Human BAT Possesses Molecular Signatures That Resemble Beige/Brite Cells.” Edited by Hironori Waki. PloS One 7 (11): e49452.

Shi, Yu, and Fanxin Long. 2017. “Hedgehog Signaling via Gli2 Prevents Obesity Induced by High-Fat Diet in Adult Mice.” eLife 6 (December). https://doi.org/10.7554/eLife.31649.

Shungin, Dmitry, Thomas W. Winkler, Damien C. Croteau-Chonka, Teresa Ferreira, Adam E. Locke, Reedik Mägi, Rona J. Strawbridge, et al. 2015. “New Genetic Loci Link Adipose and Insulin Biology to Body Fat Distribution.” Nature 518 (7538): 187–96.

Song, Tongxing, and Shihuan Kuang. 2019. “Adipocyte Dedifferentiation in Health and Diseases.” Clinical Science 133 (20): 2107–19.

Spiegelman, Bruce M., and Stephen R. Farmer. 1982. “Decreases in Tubulin and Actin Gene Expression prior to Morphological Differentiation of 3T3 Adipocytes.” Cell 29 (1): 53–60.

Spurgin, Stephen B., Lavanya Vishvanath, Karen A. Macpherson, Chelsea Hepler, Mengle Shao, and Rana K. Gupta. 2016. “Identification of Itgbl1: A Novel Regulator of Adipogenesis.” https://utswmed-ir.tdl.org/handle/2152.5/3261.

Steier, Z., McIntyre, L.L., Lutes, L.K., Huang, T.S., Robey, E.A., Yosef, N. and Streets, A., 2021. Single-cell multi-omic analysis of thymocyte development reveals NFAT as a driver of CD4/CD8 lineage commitment. bioRxiv. https://doi.org/10.1101/2021.07.12.452119.

Sun, Wenfei, Hua Dong, Miroslav Balaz, Michal Slyper, Eugene Drokhlyansky, Georgia Colleluori, Antonio Giordano, et al. 2020. “Single-Nucleus RNA-Seq Reveals a New Type of Brown Adipocyte Regulating Thermogenesis.” https://doi.org/10.1101/2020.01.20.890327.

Talebi, Ali, Jonas Dehairs, Florian Rambow, Aljosja Rogiers, David Nittner, Rita Derua, Frank Vanderhoydonc, et al. 2018. “Sustained SREBP-1-Dependent Lipogenesis as a Key Mediator of Resistance to BRAF-Targeted Therapy.” Nature Communications 9 (1): 1–11.

Tchkonia, Tamara, Nino Giorgadze, Tamar Pirtskhalava, Thomas Thomou, Matthew DePonte, Ada Koo, R. Armour Forse, et al. 2006. “Fat Depot-Specific Characteristics Are Retained in Strains Derived from Single Human Preadipocytes.” Diabetes 55 (9): 2571–78.

Tchkonia, Tamara, Yourka D. Tchoukalova, Nino Giorgadze, Tamar Pirtskhalava, Iordanes Karagiannides, R. Armour Forse, Ada Koo, et al. 2005. “Abundance of Two Human Preadipocyte Subtypes with Distinct Capacities for Replication, Adipogenesis, and Apoptosis Varies among Fat Depots.” American Journal of Physiology. Endocrinology and Metabolism 288 (1): E267–77.

Tong, Q., G. Dalgin, H. Xu, C. N. Ting, J. M. Leiden, and G. S. Hotamisligil. 2000. “Function of GATA Transcription Factors in Preadipocyte-Adipocyte Transition.” Science 290 (5489): 134–38.

Trapnell, Cole. 2015. “Defining Cell Types and States with Single-Cell Genomics.” Genome Research. https://doi.org/10.1101/gr.190595.115.

Ullah, Mujib, Michael Sittinger, and Jochen Ringe. 2013. “Extracellular Matrix of Adipogenically Differentiated Mesenchymal Stem Cells Reveals a Network of Collagen Filaments, Mostly Interwoven by Hexagonal Structural Units.” Matrix Biology: Journal of the International Society for Matrix Biology 32 (7-8): 452–65.

Urs, Sumithra, Colton Smith, Brett Campbell, Arnold M. Saxton, James Taylor, Bing Zhang, Jay Snoddy, Brynn Jones Voy, and Naima Moustaid-Moussa. 2004. “Gene Expression Profiling in Human Preadipocytes and Adipocytes by Microarray Analysis.” The Journal of Nutrition 134 (4): 762–70.

Wolock, Samuel L., Romain Lopez, and Allon M. Klein. 2019. “Scrublet: Computational Identification of Cell Doublets in Single-Cell Transcriptomic Data.” Cell Systems 8 (4): 281–91.e9.

Wu, Zeni, and Suqing Wang. 2013. “Role of Kruppel-like Transcription Factors in Adipogenesis.” Developmental Biology 373 (2): 235–43.

Xue, Ruidan, Matthew D. Lynes, Jonathan M. Dreyfuss, Farnaz Shamsi, Tim J. Schulz, Hongbin Zhang, Tian Lian Huang, et al. 2015. “Clonal Analyses and Gene Profiling Identify Genetic Biomarkers of the Thermogenic Potential of Human Brown and White Preadipocytes.” Nature Medicine 21 (7): 760–68.

Xu, Haoying, Yanlei Yang, Linyuan Fan, Luchan Deng, Junfen Fan, D. Li, Hongling Li, and Robert Chunhua Zhao. 2021. “Lnc13728 Facilitates Human Mesenchymal Stem Cell Adipogenic Differentiation via Positive Regulation of ZBED3 and Downregulation of the WNT/β-Catenin Pathway.” Stem Cell Research & Therapy 12 (1): 176.

Yamamoto, Ken-Ichi, Masakiyo Sakaguchi, Reinhold J. Medina, Aya Niida, Yoshihiko Sakaguchi, Masahiro Miyazaki, Ken Kataoka, and Nam-Ho Huh. 2010. “Transcriptional Regulation of a Brown Adipocyte-Specific Gene, UCP1, by KLF11 and KLF15.” Biochemical and Biophysical Research Communications 400 (1): 175–80.

Yang, Wulin, Xiangxiang Guo, Shermaine Thein, Feng Xu, Shigeki Sugii, Peter W. Baas, George K. Radda, and Weiping Han. 2013. “Regulation of Adipogenesis by Cytoskeleton Remodelling Is Facilitated by Acetyltransferase MEC-17-Dependent Acetylation of α-Tubulin.” Biochemical Journal 449 (3): 606–12.

Yu, Jinhai, and Peng Li. 2017. “The Size Matters: Regulation of Lipid Storage by Lipid Droplet Dynamics.” Science China. Life Sciences 60 (1): 46–56.

Zhou, Qiuzhong, Zhenzhen Fu, Yingyun Gong, Veerabrahma Pratap Seshachalam, Jia Li, Yizhe Ma, Hui Liang, et al. 2020. “Metabolic Health Status Contributes to Transcriptome Alternation in Human Visceral Adipose Tissue During Obesity.” Obesity 28 (11): 2153–62.

