## Supplemental Information for "Mapping the temporal transcriptional landscape of human white and brown adipogenesis using single-nuclei RNA-seq"

### Note S1: Pseudotemporal ordering of differentiating white and brown preadipocytes into mature adipocytes

After integration of snRNA-seq datasets from distinct time-points, a coarse cellular ordering was obtained for both white and brown fat development, with day-0 nuclei on one end, and day-20 nuclei on the other. In order to further refine this cellular-ordering for a higher resolution, we utilized the idea of pseudo-temporal analysis. Typically, pseudo-temporal analysis is performed using advanced bioinformatic algorithms that require prior information on the starting cell/cluster for ordering. Examples include Slingshot, which was identified as one of the most robust tools for pseudotemporal ordering by recent benchmarking investigations (Saelens et al., 2019). However, Slingshot does not take into account prior biological information for ordering cells (Street et al., 2018). Hence, multiple researchers have rather relied on expression values of biologically relevant genes (or highly variable genes) as proxy for cell-ordering (Cuomo et al., 2020; Kouno et al., 2013; Tran and Bader, 2020; Zeng et al., 2017). For adipogenesis, a list of such biologically relevant genes can be found as a molecular signature in the MSigDB (Hallmark_Adipogenesis, (Liberzon et al., 2011; Subramanian et al., 2005), and combined expression of these genes (or a signature “score”) could be used as a proxy to order cells. However, most genes which are part of the Hallmark_Adipogenesis signature are identified via transcriptomic enrichment analysis in mature adipocytes (terminal state), as compared to preadipocytes (initial state). Consequently, expression of such genes loses resolution for ordering cells that are in middle stages of adipogenesis. Therefore, for ordering nuclei in our dataset, we developed a strategy which utilizes expression of genes that are monotonically increasing in expression with cellular differentiation, thereby providing a high dynamic range as well as pseudo-temporal resolution. Such monotonically increasing genes were identified via a consensus of Slingshot and Hallmark_Adipogenesis ordering.

Specifically, differentiating white and brown nuclei were first ordered using both Slingshot, and Hallmark_Adipogenesis signature score (calculated using Vision, see Methods). Then, dynamically regulated genes were identified for each methodology (see Methods) and clustered based on their expression profiles. Genes that were monotonically increasing in both ordering strategies were then defined as a custom signature. Different signatures were defined specific to white and brown adipogenesis. Finally, each nuclei was assigned a score based on the expression of genes constituting these custom-defined signatures (using Vision), and this score was used as a proxy for pseudotime.

**White Signature:** *ADM	APOE	FABP5	MDH1	DCXR	AOC3	PLIN2	MME	BTG1	CD36	PDK4	FKBP5	SOX5	BCL6	ZBED3	SIK2	COL4A1	COL4A2	RPLP2	LMO4	AL845331.2	BMS1P14	SORT1	ACER3	HK2	ITGA7	SLC7A6	FZD4	TMEM135	PLA2G16	KCNIP2-AS1	KCNIP2	TNS1	DECR1	PPARGC1A	PSMA1	G0S2	PNPLA2	FABP4	MLXIPL	AQP7	ACADVL	PALMD	CYB5A	PDE3B	SAT1	ACACB	CSAD	MALAT1	DOCK11	PPARG	FOXO1	CALCRL	CHCHD10	ACO2	TOB2*

**Brown Signature:** *CYB5A	COL4A1	CIDEC	CD36	AC002066.1	CAV2	TMEM164	ACSL5	GIPR	LBP	NAMPT	PDK4	ACSL4	FADS1	ACACA	ME1	GHR	DDIT4	LPCAT3	SREBF1	NR1H3	ABCA1	SOX5	NRCAM	PDE3B	PSMA1	BCL6	DECR1	SIK2	PLIN4	PALMD	ACSL1	SAT1	TMEM135	SCD	FABP4	PNPLA2	ACACB	CSAD	PPARG	ELOVL5	TLE1	ACER3	UVRAG	LINC01239	PTK2B	MAPK10	SOS1	RHOBTB3	FKBP5	ZBTB16	EBF1	GBE1	AKR1C2*

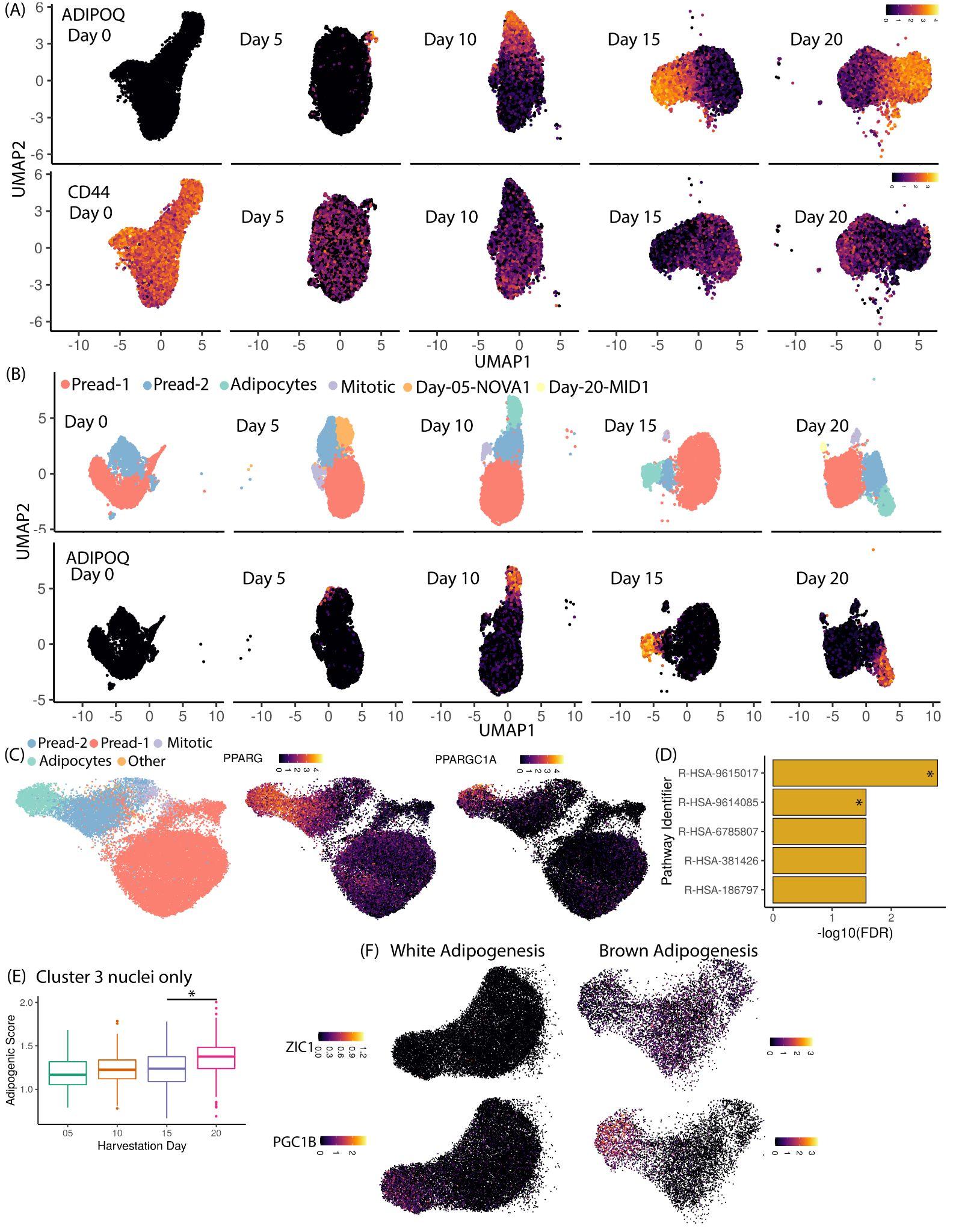

**Fig. S1 Integration of white and brown adipogenesis snRNA-seq dataset (A)** Expression of mature adipocyte marker gene *ADIPOQ* (top panel) and preadipocyte marker gene *CD44* (bottom panel) in white adipogenesis snRNA-seq datasets isolated at different time points **(B)** Unsupervised clustering of brown adipogenesis dataset (top panel) and expression of gene *ADIPOQ* (bottom panel) at different time-points **(C)** (left most panel) Integration of brown adipogenesis dataset using scvi-tools as colored by cluster assignment in panel (B) Cell types identified exclusively in one dataset (Day-05-NOVA1 and Day-20-MID1) were marked as Other. Expression of genes *PPARG* (middle panel) and *PGC1A* (right most panel) in integrated brown adipogenesis dataset **(D)** Top reactome pathways identified in the non-adipogenic response in brown dataset. R-HSA-9615017 corresponds to FOXO-mediated transcription of oxidative stress, metabolic and neuronal genes reactome pathway and R-HSA-9614085 corresponds to FOXO-mediated transcription reactome pathway. The two hits related to FOXO’s transcriptional activity are star marked **(E)** Adipogenic signature score for cluster 3 in Fig. 1H distributed over different days of harvestation **(F)** Expression of genes *ZIC1* (top panel) and *PGC1B* (bottom panel) in white and brown adipogenesis dataset

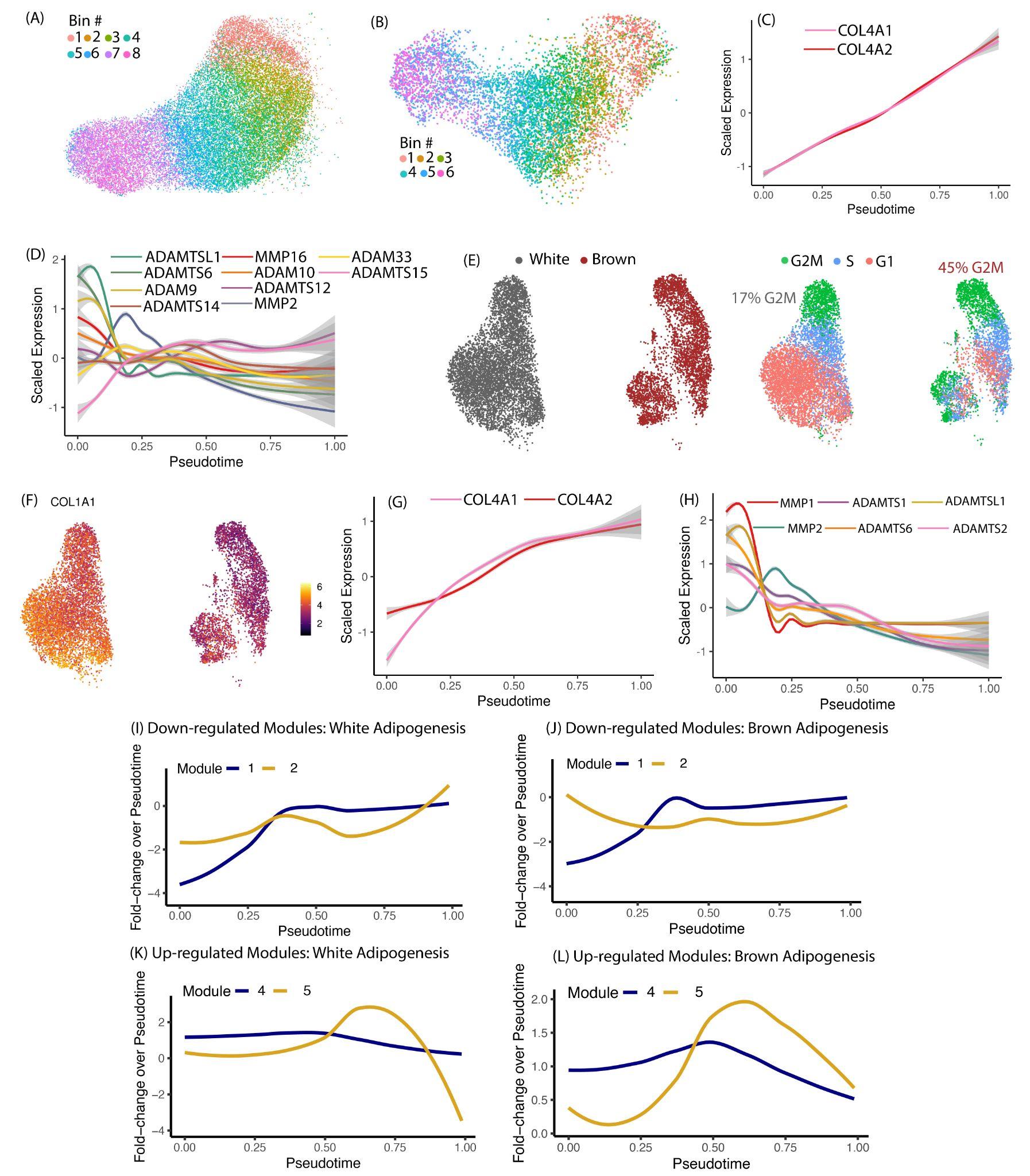

**Fig. S2 Pseudotemporal ordering of differentiating white and brown preadipocytes (A) and (B)** Pseudotemporal bins identified for differentiating white (A) and brown (B) preadipocytes. Bins were identified to have a similar number of nuclei in each bin. **(C)** Expression dynamics of *COL4A1* and *COL4A2* during white adipogenesis **(D)** Expression dynamics of metalloproteases during white adipogenesis **(E)** UMAP visualization of day-0 white and brown preadipocytes colored by lineage (left panel) and cell-cycle phase (right panel). The numbers in the right panel indicate the percent of cells in the proliferative G2M phase (F) Expression of *COL1A1* in day-0 white and brown preadipocytes **(G)** Expression dynamics of *COL4A1* and *COL4A2* during brown adipogenesis **(H)** Expression dynamics of metalloproteases during brown adipogenesis **(I and J)** Fold-change over pseudotime for Modules 1 and 2 in white (I) and brown (J) adipogenesis **(K and L)** Fold-change over pseudotime for Modules 4 and 5 in white (K) and brown (L) adipogenesis

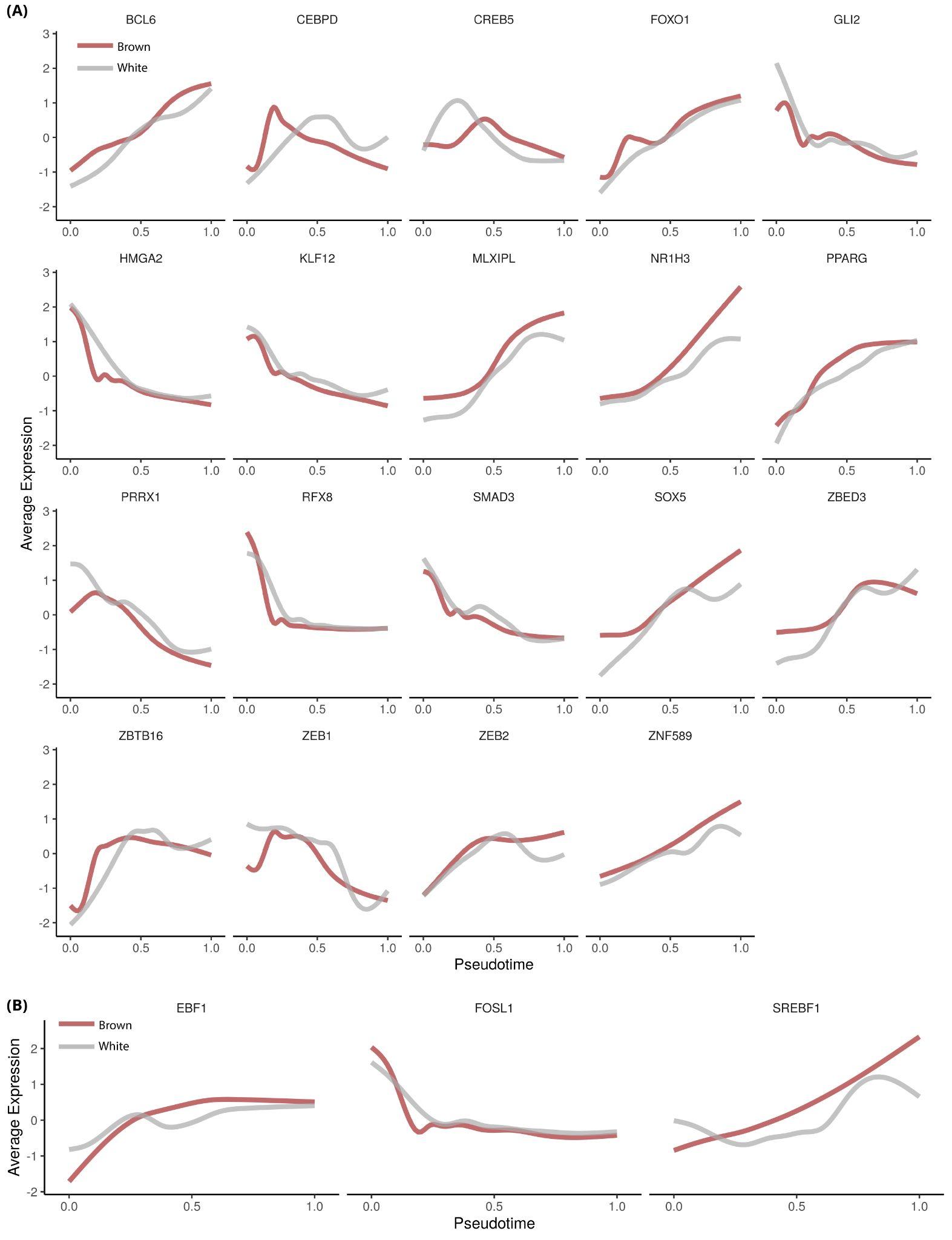

**Fig. S3 Expression dynamics of TFs in white and brown adipogenesis (A)** Dynamics of TFs temporally regulated in both white and brown adipogenesis dataset. In both datasets, all highlighted TFs had a logFC > 1 when comparing their enrichment in pseudo-temporal bins with maximum expression vs pseudo-temporal bins with minimum expression **(B)** Dynamics of TFs exclusively regulated in brown adipogenesis. For all three TFs, only brown adipogenesis dataset had logFC > 1 when comparing their enrichment in pseudo-temporal bins with maximum expression vs pseudo-temporal bins with minimum expression

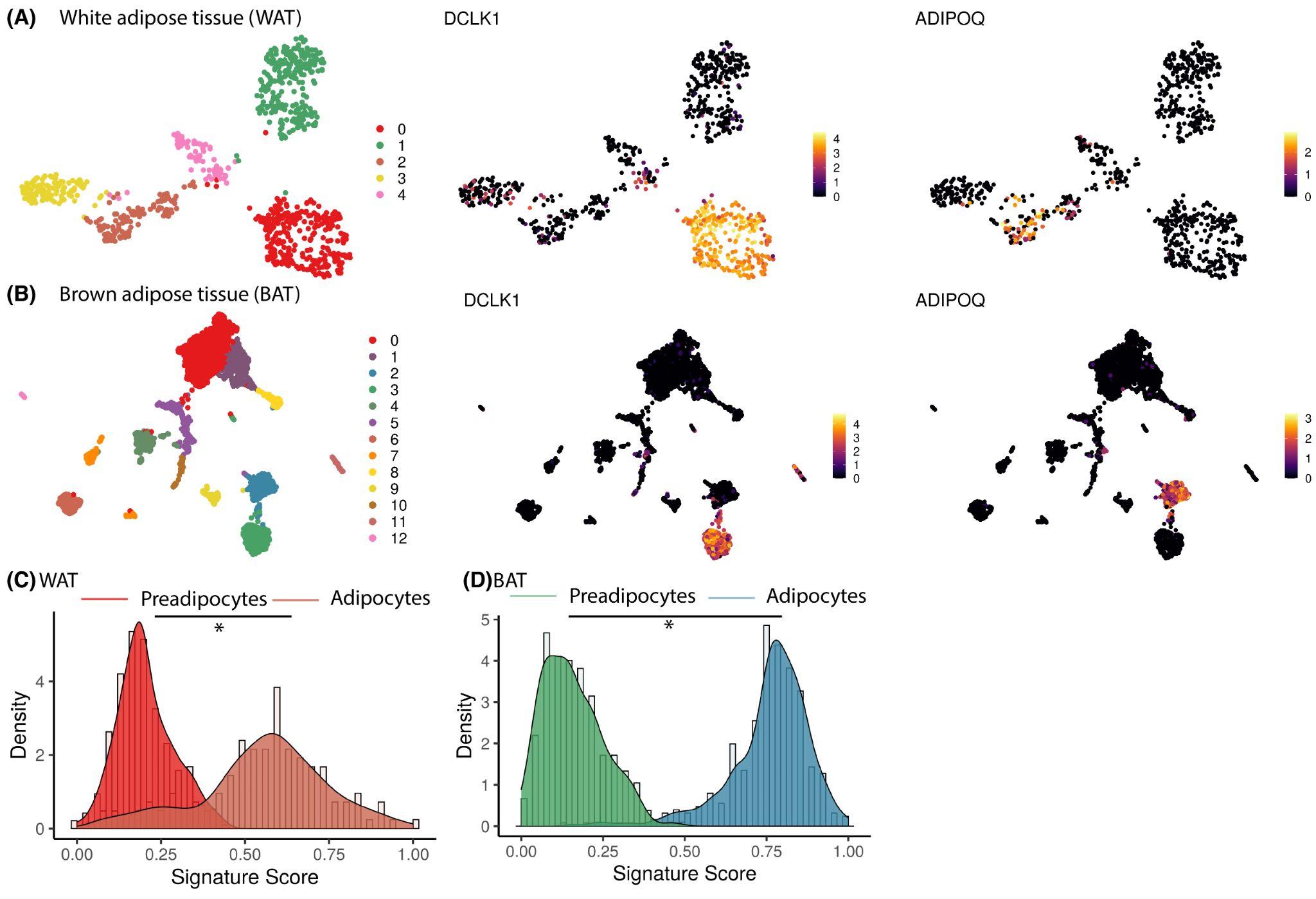

**Fig. S4 Utilization of adipogenic gene signatures for quantifying differences in cell maturation state (A)** UMAP of WAT snRNA-seq dataset colored by clusters (left panel), *DCLK1* expression (middle panel) and *ADIPOQ* expression (right panel) **(B)** Same plots as (A) but for BAT snRNA-seq dataset **(C) and (D)** Distribution of adipogenic gene signature scores between preadipocytes and mature adipocytes identified in (C) WAT and (D) BAT. Scores were scaled to vary between 0 to 1

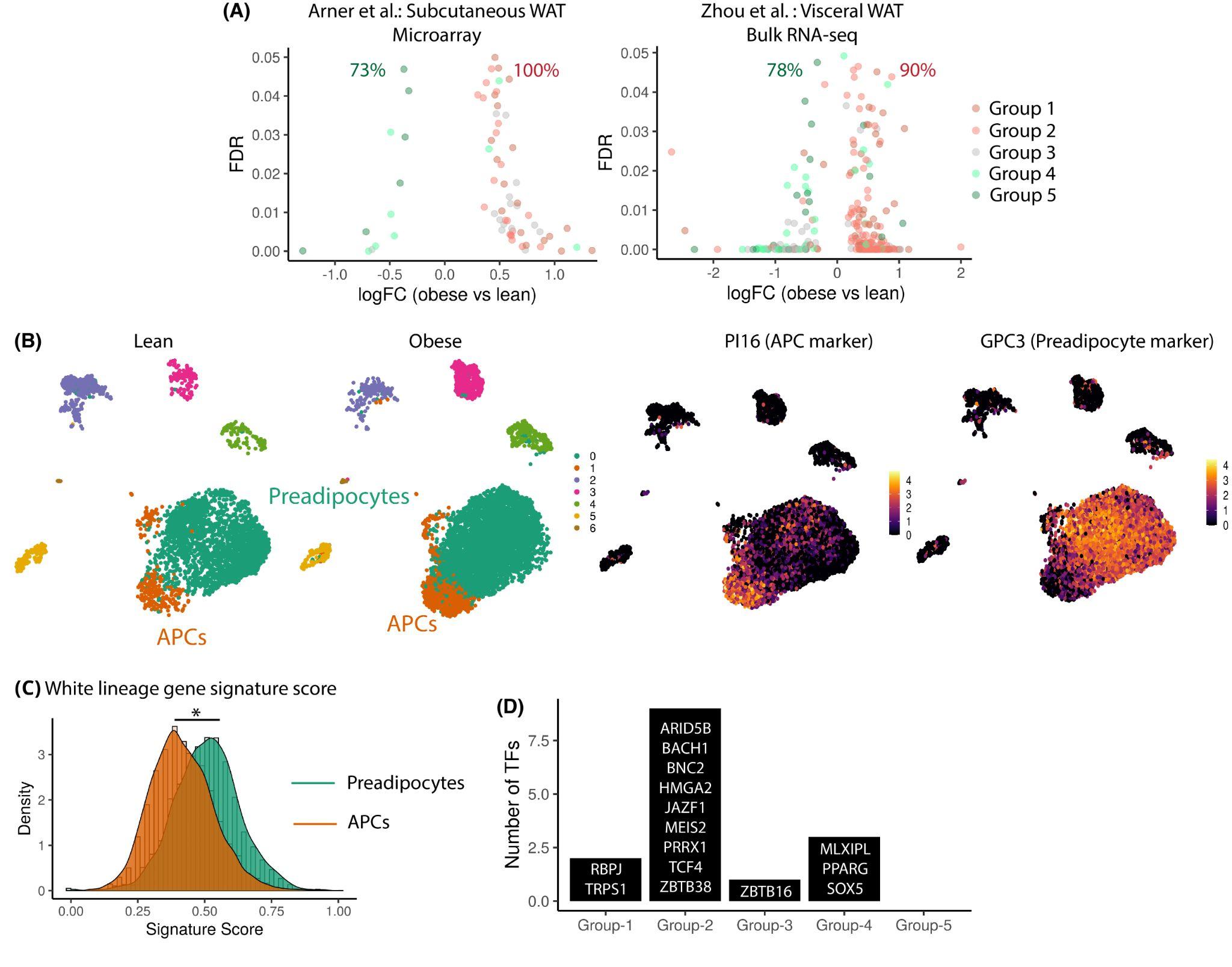

**Fig. S5 Implication of dynamically regulated genes in Obesity (A)** Volcano plot of dynamically regulated genes DE between lean vs obese bulk RNA-seq samples from two different studies. Each dot is a gene colored by its original gene module annotation. The number in green indicates percent of total DE genes in Group 4 and 5 that are enriched in lean samples. The number in red indicates percent of total DE genes in Group 1 and 2 that are enriched in obese samples **(B)** UMAP visualisation of human abdominal WAT scRNA-seq dataset colored by (left panel) clusters stratified by metabolic phenotype (middle panel) *PI16* expression (right panel) *GPC3* expression **(C)** Distribution of gene signature score between Preadipocytes and APCs identified in the same dataset as (B) **(D)** Distribution of TFs commonly identified as temporally regulated in white adipogenesis and as having SNPs in GWAS studies, distributed by their gene module annotation

### **REFERENCES**

Cuomo ASE, Seaton DD, McCarthy DJ, Martinez I, Bonder MJ, Garcia-Bernardo J, Amatya S, Madrigal P, Isaacson A, Buettner F, Knights A, Natarajan KN, HipSci Consortium, Vallier L, Marioni JC, Chhatriwala M, Stegle O. 2020. Single-cell RNA-sequencing of differentiating iPS cells reveals dynamic genetic effects on gene expression. *Nat Commun* **11**:810.

Kouno T, de Hoon M, Mar JC, Tomaru Y, Kawano M, Carninci P, Suzuki H, Hayashizaki Y, Shin JW. 2013. Temporal dynamics and transcriptional control using single-cell gene expression analysis. *Genome Biol* **14**:R118.

Liberzon A, Subramanian A, Pinchback R, Thorvaldsdóttir H, Tamayo P, Mesirov JP. 2011. Molecular signatures database (MSigDB) 3.0. *Bioinformatics* **27**:1739–1740.

Saelens W, Cannoodt R, Todorov H, Saeys Y. 2019. A comparison of single-cell trajectory inference methods. *Nat Biotechnol* **37**:547–554.

Street K, Risso D, Fletcher RB, Das D, Ngai J, Yosef N, Purdom E, Dudoit S. 2018. Slingshot: Cell lineage and pseudotime inference for single-cell transcriptomics. *BMC Genomics* **19**:477.

Subramanian A, Tamayo P, Mootha VK, Mukherjee S, Ebert BL, Gillette MA, Paulovich A, Pomeroy SL, Golub TR, Lander ES, Mesirov JP. 2005. Gene set enrichment analysis: a knowledge-based approach for interpreting genome-wide expression profiles. *Proc Natl Acad Sci U S A* **102**:15545–15550.

Tran TN, Bader GD. 2020. Tempora: Cell trajectory inference using time-series single-cell RNA sequencing data. *PLoS Comput Biol* **16**:e1008205.

Zeng C, Mulas F, Sui Y, Guan T, Miller N, Tan Y, Liu F, Jin W, Carrano AC, Huising MO, Shirihai OS, Yeo GW, Sander M. 2017. Pseudotemporal Ordering of Single Cells Reveals Metabolic Control of Postnatal β Cell Proliferation. *Cell Metab* **25**:1160–1175.e11.
